# Synthesizing Signaling Pathways from Temporal Phosphoproteomic Data

**DOI:** 10.1101/209676

**Authors:** Ali Sinan Köksal, Kirsten Beck, Dylan R. Cronin, Aaron McKenna, Nathan D. Camp, Saurabh Srivastava, Matthew E. MacGilvray, Rastislav Bodík, Alejandro Wolf-Yadlin, Ernest Fraenkel, Jasmin Fisher, Anthony Gitter

## Abstract

Advances in proteomics reveal that pathway databases fail to capture the majority of cellular signaling activity. Our mass spectrometry study of the dynamic epidermal growth factor (EGF) response demonstrates that over 89% of significantly (de)phosphorylated proteins are excluded from individual EGF signaling maps, and 63% are absent from all annotated pathways. We present a computational method, the Temporal Pathway Synthesizer (TPS), to discover missing pathway elements by modeling temporal phosphoproteomic data. TPS uses constraint solving to exhaustively explore all possible structures for a signaling pathway, eliminating structures that are inconsistent with protein-protein interactions or the observed phosphorylation event timing. Applied to our EGF response data, TPS connects 83% of the responding proteins to receptors and signaling proteins in EGF pathway maps. Inhibiting predicted active kinases supports the TPS pathway model. The TPS algorithm is broadly applicable and also recovers an accurate model of the yeast osmotic stress response.

## Introduction

High-throughput proteomic assays have illuminated the amazing breadth and complexity of the signal transduction pathways that cells employ to respond to extracellular cues. In addition to quantifying protein abundance, these technologies are now routinely used to quantify protein post-translational modifications (PTMs). Mass spectrometry, in particular, offers a broad view of PTMs, quantifying various modifications such as phosphorylation, ubiquitination, acetylation, and methylation (Choudhary and Mann, 2010). In contrast to microwestern arrays (Ciaccio et al., 2010), reverse phase protein arrays (Paweletz et al., 2001), mass cytometry (Bendall et al., 2011), and other high-throughput antibody-based assays, mass spectrometry is not restricted to a predefined list of proteins and can detect tens of thousands of phosphopeptides (Sharma et al., 2014). Here we show how to discover new facets of signaling cascades from complex proteomic data by integrating observed PTMs with existing knowledge of protein interactions.

Many gaps persist in our understanding of phosphorylation signaling cascades. For example, our mass spectrometry experiments show that nearly all proteins that are significantly (de)phosphorylated when the epidermal growth factor receptor (EGFR) is stimulated are absent from EGFR pathway maps. The low overlap is consistent with previous temporal phosphoproteomic studies of mammalian signaling (Cao et al., 2012; D’Souza et al., 2014; Humphrey et al., 2015). Discordance between mass spectrometry studies and pathway databases is partly caused by extensive crosstalk among pathways (Bauer-Mehren et al., 2009) and context-specific interactions (Hill et al., 2017). In addition, protein abundance varies greatly among human cells and tissues (Kim et al., 2014), and interactions from a pathway database are irrelevant when the proteins involved are not expressed. Moreover, perturbations and disease can rewire signaling pathways (Pawson and Warner, 2007).

Network inference algorithms can explain the phosphorylation events that lie outside of canonical pathways and complement existing manually curated pathway maps. Specialized algorithms model time series data, which contain information about the ordering of phosphorylation changes and can support causal instead of correlative modeling (Bar-Joseph et al., 2012). Temporal protein signaling information can be used to reconstruct more accurate and complete networks than a single static snapshot of the phosphoproteome.

A complementary challenge to interpreting off-pathway phosphorylation is that the cellular stimulus response includes mechanisms that are not captured in phosphoproteomic datasets. There is an interplay between phosphorylation changes and other integral parts of signaling cascades because phosphorylation can affect protein stability, subcellular localization, and recognition of interaction partners (Newman et al., 2014). Ubiquitination and other PTMs are not measured in phosphoproteomic studies, and not all phosphorylated proteins are detected by mass spectrometry. Additional information is required to infer comprehensive signaling cascades that include non-differentially phosphorylated proteins.

Protein-protein interaction (PPI) networks can be used for this purpose by identifying the chain of interactions that connect observed phosphorylation events. For example, MAP2K1 phosphorylation is not detected in our EGF response data, but our approach uses PPI to correctly determine that it is the kinase that controls MAPK1 and MAPK3 phosphorylation.

We present the Temporal Pathway Synthesizer, a method to assemble temporal phosphoproteomic data into signaling pathways that extend far beyond existing canonical maps. "Synthesizer" refers to our application of computational program synthesis techniques (Manna and Waldinger, 1980; Solar-Lezama et al., 2005) to produce pathway models from experimental data (Fisher et al., 2014), not synthetic biology (Benner and Sismour, 2005). TPS overcomes both of the aforementioned challenges in interpreting phosphoproteomic data: modeling signaling events that are not captured by pathway databases and including non-phosphorylated proteins in the predicted pathway structures. The TPS workflow consists of multiple steps (Figure 1).

**Figure 1.**
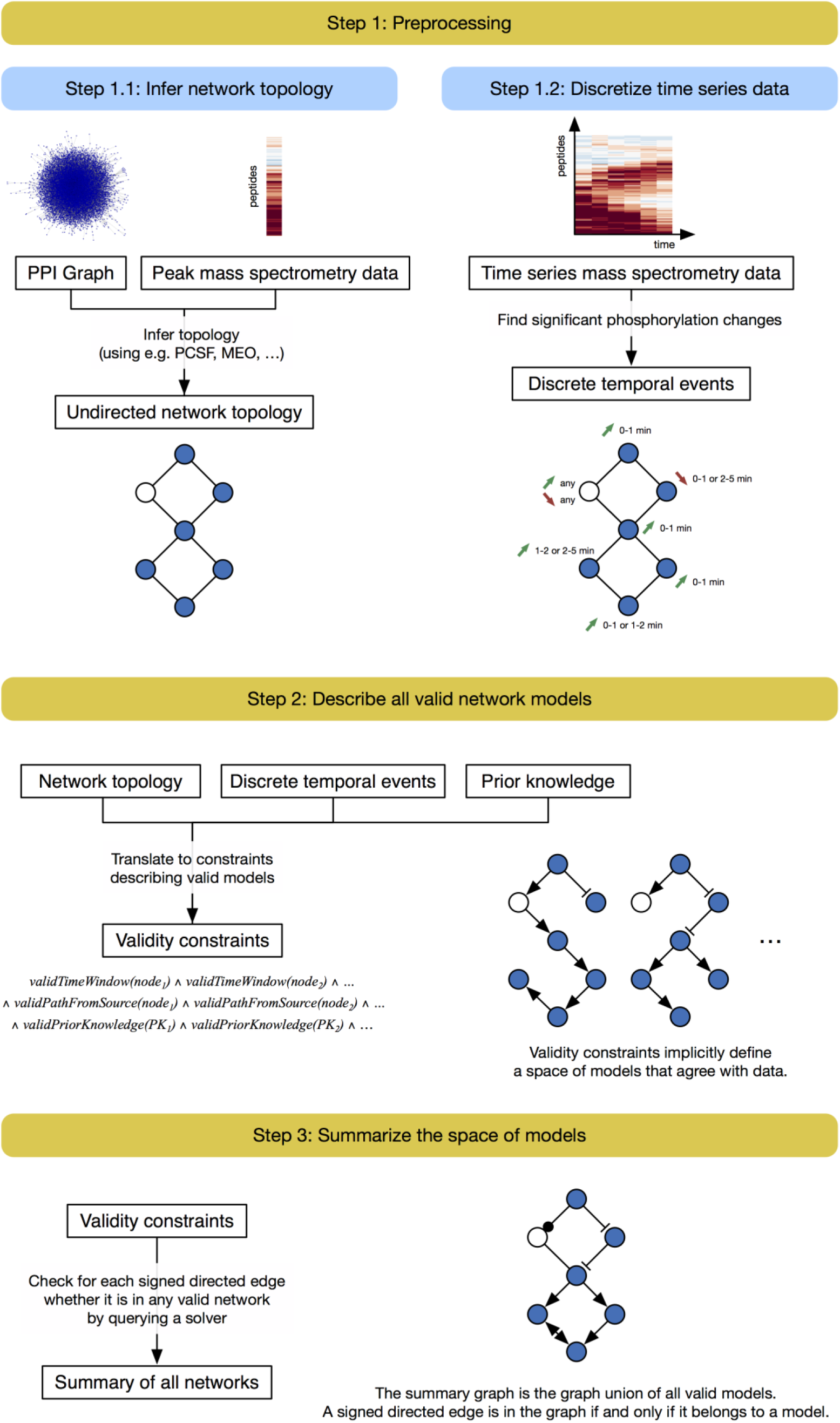
TPS workflow. First, the PPI graph is combined with the phosphorylation data to obtain a condition-specific network (Step 1.1). Algorithms used in this step do not model the temporal information. Separately, the time series data are converted into discrete timed signaling events (Step 1.2). TPS then defines a space of models that agree with the data by transforming the timed events, undirected network topology, and prior knowledge (kinase-substrate interaction directions in this study) into a set of constraints (Step 2). Our system summarizes a huge solution space by computing the union of all signed directed graph models that satisfy the given constraints (Step 3). The final pathway model predicts how a subset of generic physical protein interactions coordinate to respond to a specific stimulus in a particular cellular context.

In the first step, TPS transforms a PPI graph into a condition-specific network by using mass spectrometry data to filter out irrelevant interactions. We adopt the prize-collecting Steiner forest (PCSF) (Tuncbag et al., 2013) network algorithm to connect differentially phosphorylated proteins through high-confidence paths that may include non-phosphorylated proteins. Like nearly all existing network algorithms, PCSF cannot use temporal information.

In the second step, TPS finds the orientation and sign of edges in the condition-specific interaction graph based on the order of the phosphorylation events. Phosphorylation timing is modeled separately for each observed phosphorylation site on a protein. TPS systematically explores all possible pathway models, where each model is a signed, directed graph that explains how signaling messages propagate from the stimulated source protein. In the final step, TPS summarizes the valid models into a single aggregate network that explicitly tracks ambiguous predictions. Summarization gives insight into which edges must always take a unique sign and direction across the whole solution space and enables analysis of the large number of candidate models. We created an interactive visualization tool, the Temporal Pathway Visualizer (TPV), to display the summary network alongside the temporal phosphoproteomic data (Figure S1).

We use EGFR-mediated signaling as our primary model system for temporal phosphoproteomic and TPS analysis. TPS recovers a network that explains how EGF-responsive proteins are activated or inhibited via chains of physical interactions stemming from the EGF receptor. The highest-confidence TPS predictions are well-supported by prior knowledge and consistent with follow-up kinase inhibitions. In addition, we model the yeast osmotic stress response, recovering many of the core pathway components and predicting kinase targets that are supported by independent perturbation data. These insights into well-characterized human and yeast pathways exemplify the ability of TPS to produce condition-specific pathway maps.

## Results

### Quantitative time series phosphoproteomics of EGF response captures widespread signaling activity

To quantify global EGFR-mediated changes in cellular signaling in HEK-293 EGFR Flp-In (EGFR Flp-In) cells with phosphoproteomics, we used a well-established in-line two-dimensional high performance liquid chromatography separation (2D-HPLC) coupled to tandem mass spectrometry (MS/MS) (Ficarro et al., 2011; Wolf-Yadlin et al., 2006; Zhang et al., 2005). EGFR Flp-In cells have been used previously to study EGFR signaling *in vitro* (Gordus et al., 2009; Wagner et al., 2013), and we selected them for this study because they are easy to manipulate and provide full control of input signal. We know the number of receptors per cell and thus the ligand concentration necessary to achieve different levels of saturation. Most importantly, because EGFR Flp-In cells are homogeneous with respect to EGFR expression, this system ensures high reproducibility between replicates and minimizes effects of heterogeneous receptor expression between different samples and time points.

After EGF stimulation for 0, 2, 4, 8, 16, 32, 64, or 128 min, cells were lysed and proteins were extracted, denatured, alkylated and trypsin digested (Figure 2). Following digestion, the tryptic peptides were either lyophilized, stored for future use, or directly processed for mass spectrometry analysis. To quantify dynamic changes in protein phosphorylation, all peptides were isobarically labeled (Ross et al., 2004), enriched using phosphotyrosine-specific antibodies and/or immobilized metal affinity chromatography (IMAC) (Ficarro et al., 2002), and analyzed on a Velos Pro Elite mass spectrometer (Ficarro et al., 2011; Wolf-Yadlin et al., 2006; Zhang et al., 2005). Peptide sequences and relative quantification were determined using Comet (Eng et al., 2013). We collected three biological replicates with two technical replicates each.

Our study identifies 1,068 phosphorylation sites that are detected in all biological replicates (5,442 unique sites detected in at least one replicate), which were then used for network modeling in TPS (Tables S1 – S3 and Supplemental File 1). Phosphorylation intensities were well-correlated across the three biological replicates (Figures S2 and S3). Early temporal phosphoproteomic studies of EGF response covered fewer time points than our dataset (Olsen et al., 2006) or were limited to tyrosine phosphorylation (Oyama et al., 2009; Zhang et al., 2005). A more recent study (Reddy et al., 2016) complements ours, providing a high-temporal-resolution view of the early signaling dynamics but covering a smaller fraction of the overall EGFR pathway.

**Figure 2.**
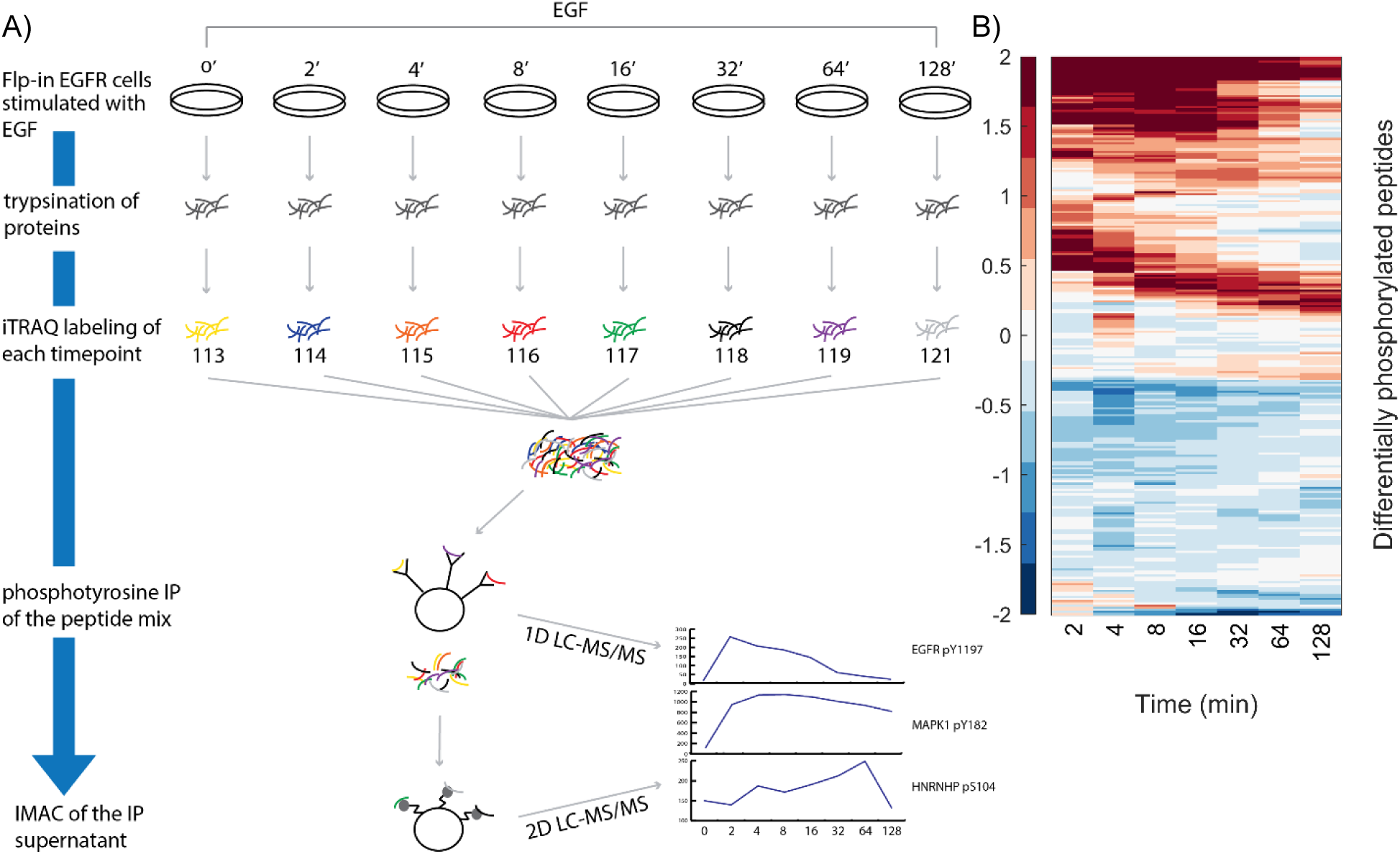
Overview of proteomics analysis. A) Cells are stimulated with EGF for 0, 2, 4, 8, 16, 32, 64, or 128 minutes and then lysed. Cellular protein content is denatured and digested. Peptides are labeled with iTRAQ and mixed. Tyrosine phosphorylated peptides are enriched by immunoprecipitation, and the flow-through is passed over immobilized metal affinity chromatography to enrich for phosphorylation events on serine and threonine. The phosphotyrosine-rich fraction is analyzed by 1D-LC-MS/MS. The more complex phospho-serine/threonine rich fraction is analyzed by 2D-LC-MS/MS. Resulting spectra are identified and quantified using Comet. B) The 263 peptides with significant temporal changes in phosphorylation exhibit distinct types of temporal behaviors (log2 fold change with respect to pre-stimulation intensity). One group of peptides is activated immediately upon stimulation, whereas others display delayed waves of phosphorylation as signals propagate.

### Reference pathway databases fail to explain phosphorylation changes

We assessed how much of the observed phosphorylation could be explained by existing pathway databases. To obtain a comprehensive view of EGFR-mediated signaling, we collected: EGFR pathway maps from six popular databases (Croft et al., 2014; Gough, 2002; Kandasamy et al., 2010; Kanehisa et al., 2012; Nishimura, 2001; Schaefer et al., 2009); a Boolean circuit representation of growth factor signaling (Layek et al., 2011); and the related but more general mitogen-activated protein kinase (MAPK) pathway from the Kyoto Encyclopedia of Genes and Genomes (KEGG). Collectively referred to as *reference pathways*, these resources reflect the diverse goals and biases of different pathway curators. BioCarta focuses on the most essential signaling events, containing only 16 proteins. Conversely, Cancer Cell Map, which is part of the NetPath resource (Kandasamy et al., 2010), seeks broader coverage. Its EGFR map contains 178 proteins, approaching the 202 proteins cataloged in a thorough EGFR review (Oda et al., 2005).

Despite the diversity of the pathway diagrams, they all fail to capture the vast majority of significant phosphorylation events triggered by EGF simulation in our system (Figures 3, S4, and S5). Among the 203 significantly differentially phosphorylated proteins, typically 5% or fewer are present in the reference pathways. The Cancer Cell Map pathway achieves the best phosphorylation coverage, but it is still only 11%. 85% of phosphorylated proteins are missing from all of the EGFR-related pathway maps (Figure S4). Additionally, most of the proteins in the EGFR pathway maps are not differentially phosphorylated (Figures 3 and S4), reflecting a combination of relevant proteins that do not undergo this particular type of PTM, phosphorylation events missed by the mass spectrometry, and interactions that are relevant in some contexts but not in EGFR Flp-In cells. The low overlaps agree with phosphoproteomic studies of other mammalian signaling pathways. Less than 10% of insulin-regulated proteins were members of a curated insulin pathway (Humphrey et al., 2015). In a study of T cell receptor signaling, only 21% of phosphorylated proteins were known to be involved in the pathway (Cao et al., 2012). Phosphosites regulated by TGF-β stimulation were not enriched for the TGF-β pathway (D’Souza et al., 2014).

Crosstalk does not explain the low coverage because most phosphorylated proteins are not present in any pathway from BioCarta, Reactome, or the Pathway Interaction Database (PID) (Figure S4). Only 37% of phosphorylated proteins are present in any pathway map. Because traditional EGFR pathway diagrams do not reflect the complex signaling observed by mass spectrometry, there is a clear need to reconstruct a context-specific representation of the underlying EGFR signaling pathway from the data.

**Figure 3.**
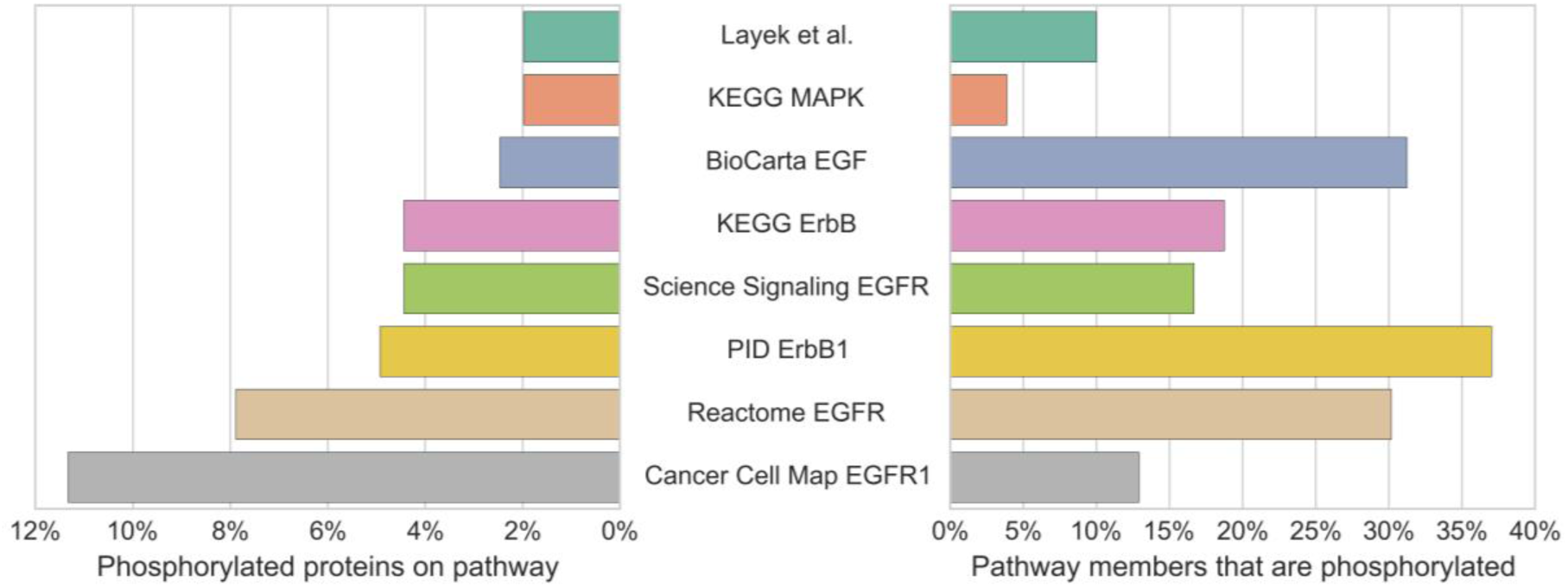
Over 95% of the significantly differentially phosphorylated proteins in response to EGF stimulation are not included in six of the reference pathways. Conversely, the majority of proteins in the reference pathways are not significantly differentially phosphorylated.

### Reconstructing the EGFR pathway with TPS explains temporal phosphorylation changes

We applied TPS to model the dynamic signaling response to EGFR stimulation in EGFR Flp-In HEK-293 cells. Our workflow consists of three major steps: (1) preprocessing the protein-protein interaction network and temporal phosphorylation data; (2) transforming temporal information, subnetwork structure, and prior knowledge into logical constraints; and (3) summarizing all valid signaling pathway models to discover interactions with unambiguous directions and/or signs (Figure 1). This process is fully described and illustrated with a simple example in Experimental Procedures.

We first discretized the time series phosphoproteomic data, using Tukey's Honest Significant Difference (HSD) test (Yandell, 1997) to determine whether a peptide exhibits a significant increase, significant decrease, or no change in phosphorylation at each post-stimulation time point. Significant phosphorylation changes can be relative to either the pre-stimulation baseline level or the previous time point. 263 peptides, corresponding to 203 proteins, significantly change at one or more time points (Figure S6). Second, we used PCSF to link the phosphorylated proteins to EGF, the source of stimulation, weighting proteins based on their HSD test significance. PCSF identifies a PPI subnetwork of 316 nodes and 422 edges (Supplemental File 2). This subnetwork comprises the interactions through which signaling messages are most likely to propagate. Third, TPS combined the discretized temporal activities of the 263 significantly changing peptides, the PCSF network, and prior knowledge (the orientation of kinase-substrate interactions) to generate a summary of all feasible pathway models (Table S4). Each type of input was translated into logical constraints, which were used to rule out pathway models that are not supported by the data.

In contrast to the reference EGFR pathway diagrams, which capture at most 11% of the differentially phosphorylated proteins, the predicted network from TPS (Figures 4 and S7 and Supplemental File 3) contains 83% of the responding proteins in its 311 nodes. Each of these proteins can be linked to the EGF stimulation with high-confidence PPI and has timing that is consistent with the temporal phosphorylation changes of all other proteins in the pathway. In addition to the phosphorylated proteins, 38 other proteins are included in the signaling pathway as hidden intermediate nodes that propagate signals via different mechanisms. Some of the differentially phosphorylated proteins may not be functional, but the TPS network provides a framework to study their role in the EGF response. The TPS pathway model includes 41 kinases and 5 phosphatases as well as adaptors and other types of proteins that coordinate with the direct phosphorylation regulators.

**Figure 4.**
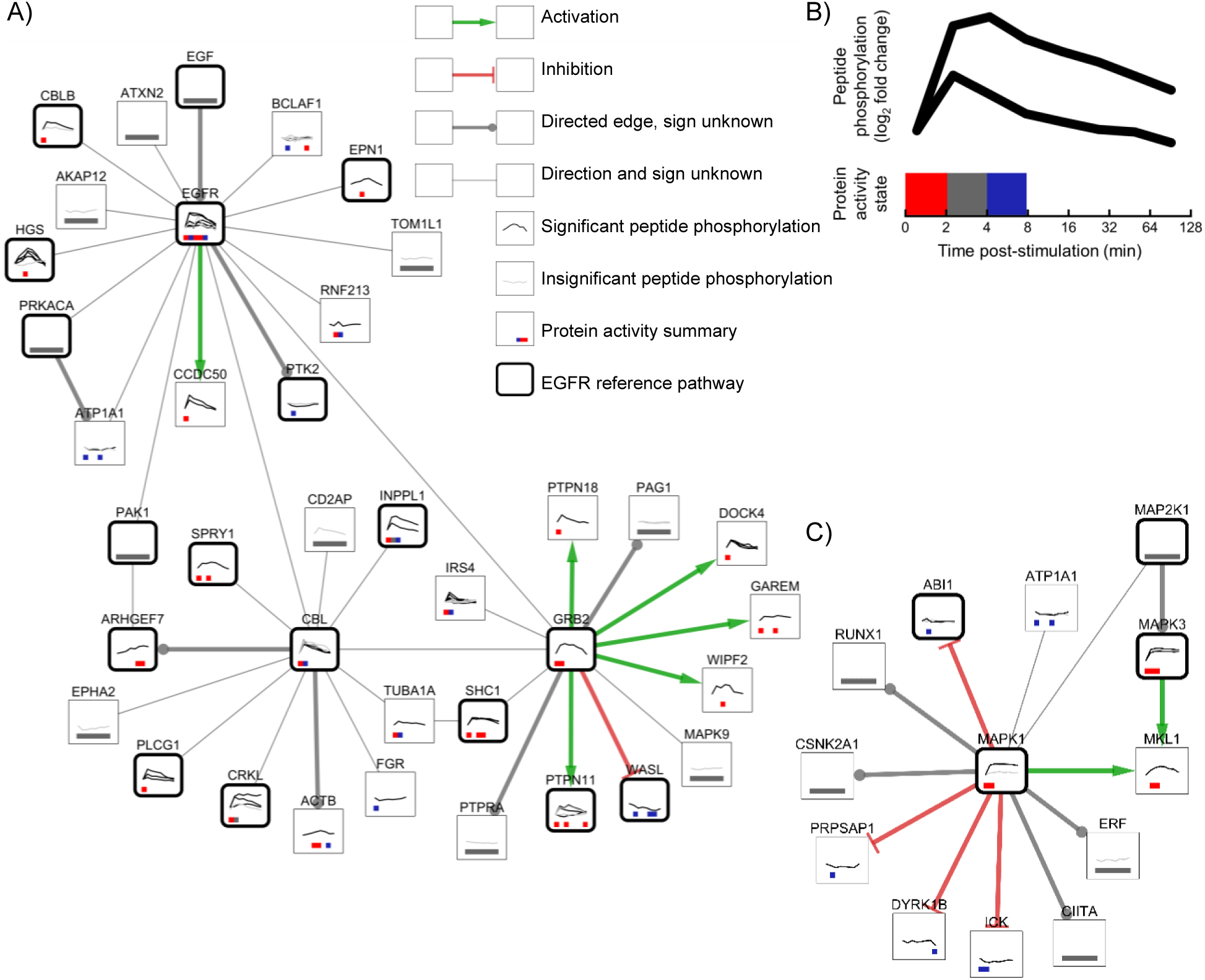
Zoomed regions of the full TPS pathway model (Figure S7). A) The EGFR subnetwork (EGFR, GRB2, CBL, and all their direct neighbors) depicts the proteins that first react to EGF stimulation. A substantial portion of the EGFR subnetwork (18 of 38 proteins) is known to be associated with EGFR signaling. Green and red edges depict activation and inhibition, respectively. Gray edges that terminate in a circle indicate that the interaction is used in the same direction in all possible pathway models, but the sign is ambiguous. Thin, undirected edges are used in different directions in different valid pathway models. Thick, rounded borders show which proteins are present in one or more reference EGFR pathways. Node annotations are detailed in panel B. B) Line graphs on each protein node show the temporal peptide phosphorylation changes relative to the pre-stimulation level on a log2 scale. Multiple lines indicate multiple observed phosphopeptides for that protein, where black lines denote statistically significant phosphorylation changes and gray lines indicate insignificant changes. Proteins without line graphs are connective Steiner nodes inferred by PCSF. Colored boxes summarize the TPS inferred activity state across peptides at each time point. Red indicates activation, blue inhibition, gray ambiguity, and white inactivity. C) The subnetwork surrounding MAPK1 and MAPK3. TPS uses the PPI network to correctly determine that MAP2K1 is the kinase that controls both MAPK1 and MAPK3 even though it is not observed in the mass spectrometry data.

Like reference pathway maps, the TPS network traces the physical protein interactions used to transmit messages from EGF to the phosphorylated proteins, including PTMs and other types of interactions. These interactions are depicted as directed, signed edges in a graph, where the sign reflects that the proteins have the same (activation) or opposite (inhibition) activity changes. The timing of the phosphorylation changes supports many possible valid interpretations, and the TPS summary tracks which edges are used in different manners in different models. Of the 413 edges in the network, 202 (49%) have a consistent direction in *all of the valid pathway models*, a very strong assertion about the confidence in these edge directions.

Thirty-eight of these directed edges have a consistent sign as well. The PPI connections, phosphorylation timing, and prior knowledge of kinase-substrate interaction direction all play distinct, important roles in reducing the number of valid pathway models (Supplemental Results, Figure S8, and Tables S5, S6, S7, and S8). The timing of protein activation and inactivation in the TPS pathway reveals a rapid spread of signaling post-stimulation (Supplemental Results and Tables S9 and S10).

Although nearly all differentially phosphorylated proteins lie outside traditional EGFR pathway representations, components of the TPS pathway reflect established EGFR relationships. Twenty-nine (11%) of the 273 phosphorylated proteins and 5 (13%) of the 38 unphosphorylated connective proteins in the TPS network are recognized as EGFR pathway members, consistent with our expectations based on the low overlaps between the significantly phosphorylated proteins and the EGFR pathway maps (Figure 3). Four sites on EGFR are immediately and significantly phosphorylated post-stimulation as are two others at later time points. Pathway models could potentially begin with EGF interacting with any of its 17 partners in the PPI network, but strong EGFR phosphorylation leads TPS to initiate all paths with the edge EGF->EGFR. Proteins directly connected to EGFR include known pathway members CBL, CBLB, EPN1, HGS, GRB2, PAK1, PRKACA, and PTK2 (Figure 4A). SHC1 is connected to EGFR via GRB2, but the direction and sign of the interaction are ambiguous due to a second parallel connection through CBL, which is also active at the two- and four-minute time points. Likewise, PLCG1 is connected via CBL, again with indeterminate direction and sign. This reflects how quickly EGFR and the other upstream pathway members are activated, suggesting that sub-minute time points may be required to unambiguously order some of the immediate connections adjacent to EGFR (Reddy et al., 2016).

Further downstream, MAP2K1 is one of several canonical EGFR pathway members that are not phosphorylated in our mass spectrometry data but are included in the pathway. Such proteins emphasize the necessity of including PPI in the analysis of the temporal phosphorylation changes because these unobserved proteins could not be recovered by any algorithm that reconstructs the pathway from the mass spectrometry data alone. MAP2K1 is correctly recognized as the direct kinase of EGFR pathway members MAPK1 (after adding new experimental constraints described below) and MAPK3 (Figure 4C). MAPK1 and MAPK3 phosphorylation levels are highly correlated and would likely be directly linked by an approach based on correlation or mutual information, but TPS correctly predicts that MAPK1 and MAPK3 correlation is due to the common upstream regulator (MAP2K1) instead. Immediately downstream of these proteins, MKL1 phosphorylation is not as strongly correlated as the two MAPKs, but TPS combines the topological constraints with the temporal information to correctly recover MAPK1->MKL1 and MAPK3->MKL1 (Muehlich et al., 2008).

### Prior evidence supports directions of EGFR pathway predictions

We find strong literature support for many of the directions that TPS predicts in the reconstructed EGFR pathway. In total, 82 of 202 interaction directions are verified in our semi-automated evaluations, and the vast majority of the remaining directions can neither be confirmed nor refuted (Table S4). The most compelling evidence comes from the EGFR reference pathways, which confirm both the edge direction and relevance to EGF stimulation response. Seven directed edges appear in one of the reference pathways, four with the predicted direction and three in complexes (Supplemental Experimental Procedures and Supplemental File 4). We expect this overlap to be low because so few significantly phosphorylated proteins are in the reference pathways (Figure 3). In addition, 78 directed edges come from the PhosphoSitePlus input data (Hornbeck et al., 2015), in which the kinase-substrate interaction direction is already known to be correct in other contexts. Nearly all of these interactions were not previously reported to be involved in EGFR signaling; only three are present in an EGFR reference pathway. We use natural language processing (NLP) software (Chen and Sharp, 2004; Hoffmann and Valencia, 2004; Poon et al., 2014) to broaden our edge direction evaluation (Supplemental Experimental Procedures). The NLP tools confirm the directions of fifteen predicted interactions and contradict only one prediction, but a manual literature review supports that prediction as well (Supplemental Results and Table S4).

### Inhibitor-induced phosphorylation changes support novel EGFR pathway predictions

To further assess components of the TPS network, we analyzed several interactions that are not present in EGFR pathway databases. These proteins are already known to physically interact. The novelty of the TPS predictions is the interactions’ relevance to the EGF response. We prioritized interactions that extend the existing EGFR pathways. Specifically, we focused on edges for which the direction or sign were predicted confidently and one of the two proteins is a member of an EGF response reference pathway (Supplemental Results). For each interaction, we inhibited the predicted upstream protein and measured the effect on the predicted target’s phosphorylation using Western blotting. From our list of ten candidate interactions (Table S11), we selected the three edges for which the antibodies reliably produced clean and quantifiable bands at the right molecular weight: MAPK1-ATP1A1, ABL2->CRK, and AKT1->ZYX (Zyxin) (Supplemental Results and Figures 4C and 5). The inhibitors used to inhibit the upstream proteins were SCH772984 for MAPK1, Dasatinib for ABL2 and MK-2296 for AKT1. After serum-starvation, the cells were treated with an inhibitor for one hour and then stimulated with EGF. We collected data at two time points (denoted short and long, see Figure 6) based on the timing of the phosphorylation events in our mass spectrometry data. Lysates were then assayed by Western blot to quantify the level of phosphorylation of the downstream protein.

**Figure 5.**
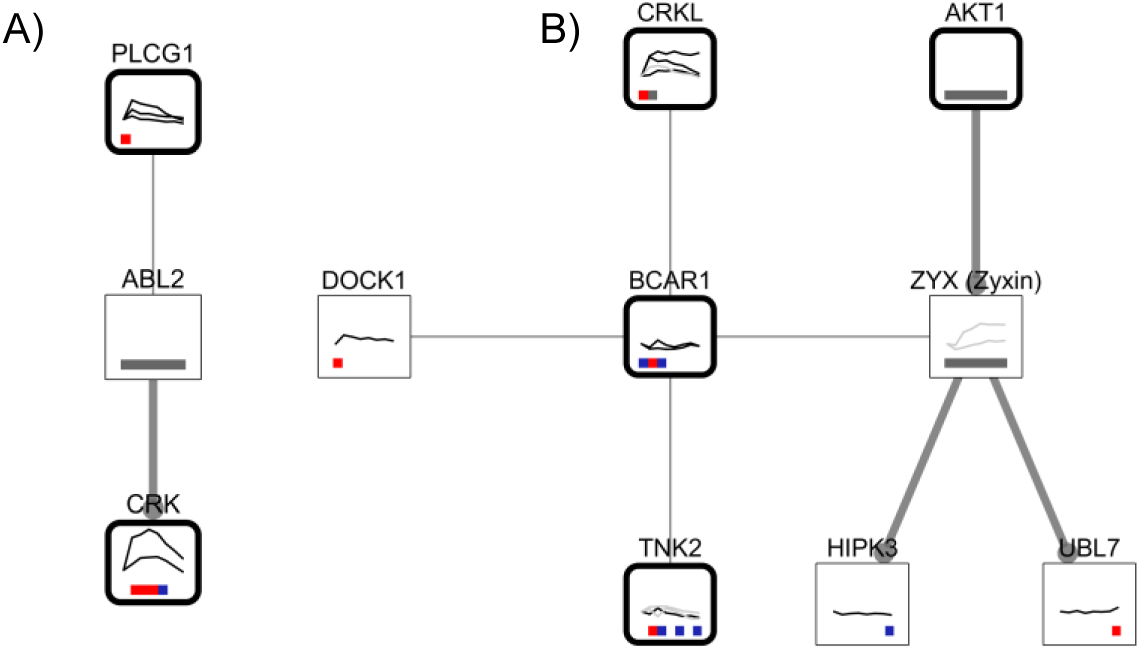
Predicted edges selected for experimental testing. The pathway context of the MAPK1 and ATP1A1 interaction is shown in Figure 4C. A) The predicted pathway context of the ABL2 to CRK interaction. B) The pathway context of the AKT1 to Zyxin interaction, which includes BCAR1, Zyxin, and all of their neighbors.

The inhibition of ABL2 decreased phosphorylation of CRK (isoform Crk-II) pY221, consistent with the TPS pathway edge (Figure 6). Inhibiting AKT1 increased phosphorylation of Zyxin. In both cases, the predicted interaction direction is supported. MAPK1 inhibition increased ATP1A1 pY10 phosphorylation. The TPS model predicted an inhibitory interaction between these proteins, but the direction was ambiguous. Our data agree with the predicted edge sign and suggest that MAPK1 is upstream of ATP1A1. These results provide independent experimental evidence that is consistent with the novel edges identified in our network analysis. However, because the inhibitors may also act upon other related kinases and the kinases’ regulation may not be direct, further experiments are required to fully validate the predictions.

**Figure 6.**
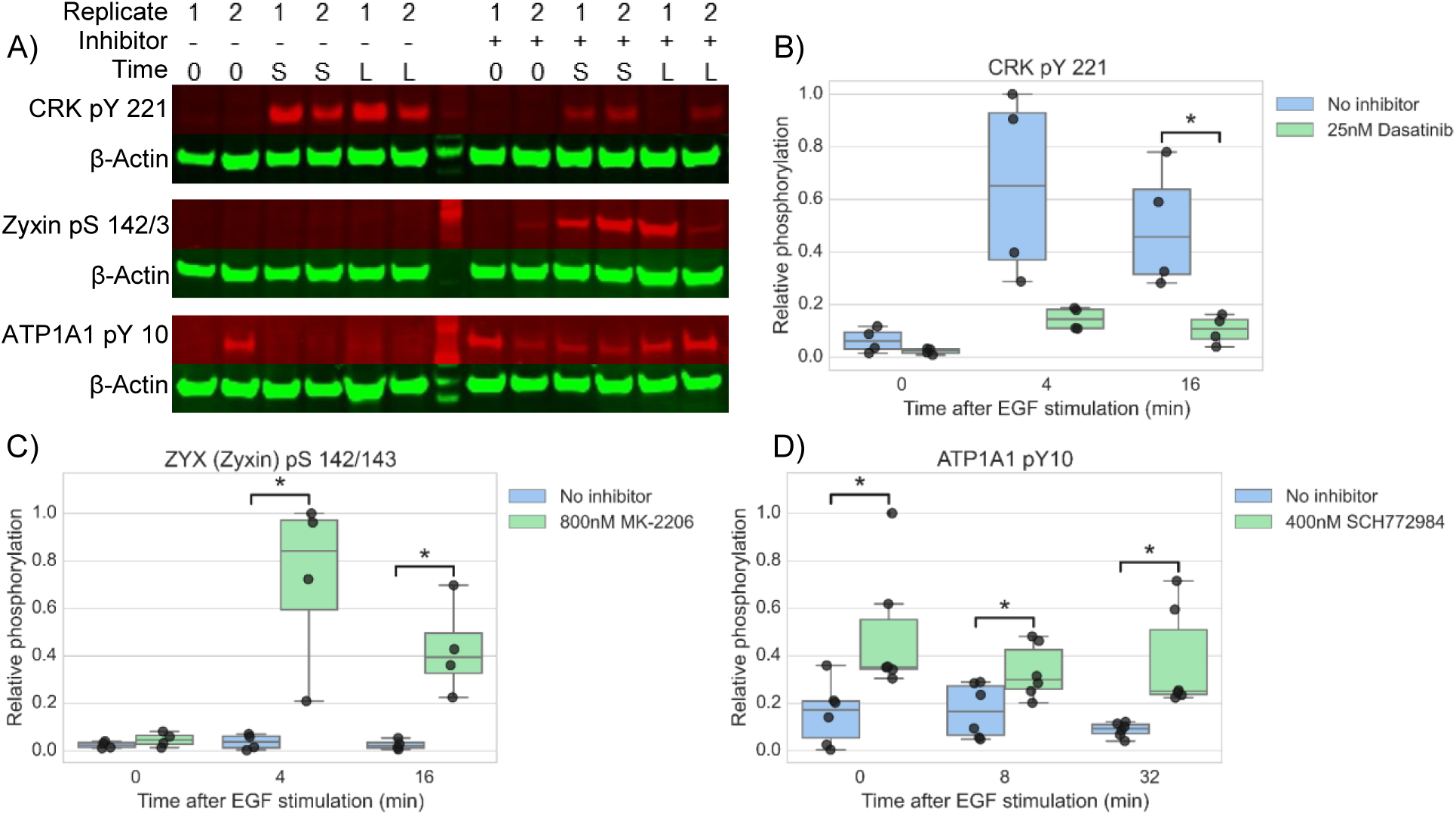
Inhibiting predicted pathway edges. A) Western blots for CRK pY 221, Zyxin pS 142/143 and ATP1A1 pY 10 in the presence and absence of small molecule inhibitors targeting their parent node (Dasatinib/ABL2, MK-2206/AKT1, and SCH772984/MAPK1, respectively). The red channel displays detection of the specific phosphorylation sites. The green channel displays detection of β-Actin (a loading control used for normalization of each specific phospho signal). Two biological replicates are shown in each Western blot. Time “0” indicates no EGF stimulation; “S” is short EGF stimulation (four or eight min), and “L” is long EGF stimulation (sixteen or thirty-two min). Absence or presence of inhibitor is shown by “−” and “+”, respectively. B) Quantification of CRK pY 221 phosphorylation (four replicates). Phosphorylation levels are relative to the maximum phosphorylation across all conditions and replicates. An asterisk denotes p < 0.05 (two-sided, unpaired, unequal variances t-test). Whiskers show 1.5 times the interquartile range. C) Quantification of Zyxin pS 142/143 phosphorylation (four replicates). D) Quantification of ATP1A1 pY 10 phosphorylation (six replicates).

### Iterative experimental and computational modeling further reduces pathway ambiguity

An important feature of TPS is its flexibility to integrate different types of constraints on pathway structure. This makes it ideal for iterative modeling because computational hypotheses that are experimentally confirmed or refuted can be fed back into TPS. Based on the results from the Western blots, we added a new constraint: MAPK1 inhibits ATP1A1. We then ran TPS again, requiring pathways to be consistent with the new constraint and all previous constraints. After restricting the pathway structure, TPS correctly infers that MAP2K1 is directly upstream of MAPK1 yielding a more precise and accurate pathway. Without the Western blot-derived constraint, the direction of the MAP2K1-MAPK1 interaction was ambiguous due to the possibility that ATP1A1, rather than MAP2K1, controls MAPK1 phosphorylation changes (Figure 4C). Other types of experimental corroboration can be similarly applied to iteratively improve the predictive power of TPS.

### TPS makes network predictions not captured by alternative approaches

We compared TPS to two existing methods that combine PPI networks and time series data and a third that uses only time series data (Supplemental Experimental Procedures). The dynamic Bayesian network (DBN) (Hill et al., 2012) infers posterior peptide-peptide interaction probabilities from time series data and network priors. TimeXNet (Patil and Nakai, 2014; Patil et al., 2013) formulates pathway prediction as a network flow problem. FunChisq (Zhang and Song, 2013) uses an adapted chi-square test to detect directed relationships between phosphorylated proteins. Comparing the four predicted pathways demonstrates the impact of the diverse algorithmic strategies. Almost all of the protein-protein edges are unique to a single method, and no edges are predicted by all four methods (Figure S9A). Despite greater overlap among the predicted nodes (Figure S9B), the four pathways are divergent. The TPS constraints allow it to recover different pathway relationships from those identified by the existing methods.

Because most of the differentially phosphorylated proteins are not members of any reference pathway, these pathways cannot be used to assess the overall quality of the predictions. However, it is still informative to compare the methods on the small fraction of predictions that are covered in the databases. The TimeXNet pathway, the largest of the three predicted networks, generally captures the most reference pathway interactions when ignoring edge direction and sign (Table S12). However, a closer examination that accounts for the predicted interaction direction shows that TPS typically makes the fewest errors, even when controlling for the size of the predicted pathways (Table S12). TPS only asserts an edge direction when it is certain that all plausible pathway models must use an interaction in a particular direction. As in the NLP evaluation, this conservative approach leads to very accurate direction predictions, which is crucial for designing validation experiments. When tested against the collection of EGFR reference pathways and all pathways in the BioCarta, Reactome, and PID databases, TPS makes only one incorrect direction prediction, though many of the directed edges are not in the reference pathways and cannot be confirmed or refuted. In contrast, the DBN and TimeXNet predict more directed edges but make errors at a greater rate.

The reference pathway evaluation illuminates additional ways in which our constraint-based approach improves upon the existing methods. Unlike TPS, the DBN and FunChisq can infer only interactions between proteins observed in the phosphorylation time series data and ignore unphosphorylated nodes. Therefore, these methods miss important interactions involving proteins with no associated time series data, which TPS can detect. These include two of our experimentally-tested predictions (ABL2->CRK and AKT1->ZYX) and additional interactions with EGFR reference pathway members MAP2K1 and PAK1. Overall, the DBN predicts more edges than TPS, but fewer of those edges are in the reference pathways (Table S12). FunChisq also produces a larger network than TPS but has no directed edges that overlap with directed edges in the reference pathways. Inspection of the FunChisq network reveals that this may be because its predicted network does not resemble the topology of the reference pathways. It contains several clusters of nodes, and nodes within a cluster are predicted to interact with most other nodes in the cluster.

By making greater use of the temporal information in the time series data, TPS can detect temporal inconsistencies in TimeXNet predictions that conflict with the reference pathway directions. We identified incorrectly oriented TimeXNet predictions, such as GRB2->EGFR and GRB2->ERBB2, for which TPS detects an inconsistency between the prediction and the input data. When providing the TimeXNet output as input to TPS, TPS reports that there is no pathway structure that can activate GRB2. Consequently, the GRB2 node and the incorrect EGFR and ERBB2 interactions are removed. As an example of an invalid pathway structure, TimeXNet predicts that signal flows from LCK to CBL to GRB2. However, TPS shows that any directed path from LCK to GRB2 violates the phosphorylation timing because LCK is first differentially phosphorylated at 16 min, later than GRB2’s initial change at 4 min. TPS detects similar contradictions for all other paths to GRB2 in the TimeXNet network. Although TimeXNet also models phosphorylation timing, it cannot model the dynamics of multiple phosphosites per protein and uses a coarse division of the time series data into three bins, which can produce misleading temporal interpretations (Supplemental Results).

### TPS generates high-quality pathway models from lower temporal resolution phosphoproteomic spectrometry data

Having established that TPS produces informative signaling pathway models from our EGF stimulation phosphorylation data, we assessed whether it could attain similar performance on existing temporal phosphoproteomic datasets, which typically cover fewer proteins and time points. A foundational study by Olsen et al. (Olsen et al., 2006) examined the phosphorylation response to EGF at 0, 1, 5, 10 and 20 minutes post-stimulation. We identified 302 phosphopeptides on 203 proteins with a phosphorylation log2 fold change of at least 2 and constructed PCSF and TPS pathway models from these proteins that exhibited temporal phosphorylation changes (Supplemental Files 5 and 6). The resulting TPS model connects 280 proteins with 329 edges, and 242 edges can be assigned a unique direction in all possible pathway structures (Figure S10). Twenty-four (10%) of the 243 phosphorylated proteins and 9 (24%) of the 37 unphosphorylated connective proteins are known EGFR pathway members. We evaluated the 242 edges assigned a unique direction in the TPS model against the EGFR-specific and generic reference pathways. The median number of directed edges that matched a directed edge in a reference pathway was two, and none of the reference pathways reported an interaction that conflicted with any of the directed predictions. Overall, TPS performs well even with fewer time points and differentially phosphorylated proteins.

Although the TPS pathways produced with our phosphoproteomic data and the Olsen et al. data are of similar sizes and contain an almost identical number of proteins from the EGFR reference pathways, a closer examination reveals that their pathway structures reflect the differences in the underlying phosphorylation data. The Olsen et al. EGF stimulation study used HeLa cells, a shorter time course, and different phosphorylation quantification techniques than our approach and did not provide biological replicates. This led to substantial differences in the quantified proteins and the magnitudes of their phosphorylation changes (Figure S11), which consequently have a strong influence on the resulting network models. Only 59 proteins and 25 edges are common to both TPS models, and each TPS EGFR network captures unique parts of the reference EGFR pathways (Figure S12). Each network reflects the individual aspects of EGF signaling in the cell types and conditions being studied, though it is likely that the quantified phosphosites and generated TPS models would grow more similar with deeper phosphoproteome coverage.

### Yeast osmotic stress response model recapitulates known pathway structure and nominates candidate Rck2 and Cdc28 substrates

Although they are still not fully characterized, stress-response signaling cascades in the yeast *Saccharomyces cerevisiae* are better-understood than their human counterparts and are not subject to cell type-specific effects. Thus, we applied TPS to model the yeast osmotic stress response to assess its ability to recapitulate this frequently-studied pathway and reveal additional novel interactions. The hyperosmotic stress response is primarily controlled by the High Osmolarity Glycerol (HOG) pathway and its central mitogen-activated protein kinase Hog1, which regulates osmo-responsive transcriptome changes (Chasman et al., 2014; O’Rourke and Herskowitz, 2004; Posas et al., 2000; Rep et al., 2000), transient translational inhibition (Melamed et al., 2008; Teige et al., 2001; Uesono and Toh-e, 2002), cell cycle arrest (Clotet et al., 2006; Duch et al., 2012; Escoté et al., 2004), and metabolic changes controlling intracellular osmolyte levels (Albertyn et al., 1994; Dihazi et al., 2004; Norbeck et al., 1996). Kanshin et al. profiled the rapid response to NaCl, an osmotic stressor, measuring phosphorylation changes for 60 seconds post-stimulation at uniform 5 second intervals (Kanshin et al., 2015). They identified 1,596 phosphorylated proteins, including 1,401 dynamic phosphopeptides on 784 proteins based on their fold changes in the salt stress time series with respect to a control (Table S13). We used these data to construct a TPS pathway model of the early osmotic stress response (Supplemental Files 7 and 8).

The TPS osmotic stress pathway contains 216 proteins and 287 interactions (Figure S13). Thirty-six of these proteins (17%) have been previously annotated as osmotic stress pathway proteins (Kawakami et al., 2016). Focusing on the subset of interactions that connect known HOG pathway members reveals that many of the edges connecting them are correct as well (Figure S14A). TPS recovers the core part of the KEGG high osmolarity pathway, including the interactions Sho1->Ste50, Sho1->Cdc24, Sho1->Pbs2, Ssk2->Pbs2, and Pbs2->Hog1. In addition, it correctly places Hog1 as the direct regulator of Rck2 (Bilsland-Marchesan et al., 2000) and the essential osmotic stress response transcription factors Hot1, Msn2, and Sko1 (Capaldi et al., 2008). TPS identifies Sch9 as an additional regulator of Sko1 (Pascual-Ahuir and Proft, 2007). Following hyperosmotic shock, Hog1 is recruited to Fps1 (Lee et al., 2013), consistent with the TPS prediction. The predicted feedback from Hog1 to Ste50 is also well-supported in osmotic stress (Hao et al., 2008). Many predicted interactions that deviate from the canonical HOG pathway model can be attributed to the input phosphorylation data and background network, not the TPS algorithm (Supplemental Results).

After confirming the TPS osmotic stress model agrees well with existing models, we investigated novel candidate pathway members. Upon a shift to high osmolarity, Hog1 phosphorylates and activates the kinase Rck2 (Bilsland-Marchesan et al., 2000), which then phosphorylates the translation elongation factor 2 (Eft2), temporarily inhibiting translation (Teige et al., 2001). The TPS model captured the cascade Hog1->Rck2->Eft2 and predicted additional Rck2 targets (Figure S14B). To test these predictions, we compared them to a recent phosphoproteomic study of an *RCK2* mutant subjected to osmotic stress (Romanov et al., 2017). All four proteins that TPS predicts are activated by Rck2 have defective phosphorylation on at least one phosphosite in *rck2*Δ five minutes after osmotic insult (Romanov et al., 2017). Thus, Rck2 likely directly phosphorylates Fpk1, Pik1, Rod1, and YLR257W upon osmotic stress, as TPS predicts. In addition to the four activated substrates, TPS predicts that Rck2 regulates YHR131C. In this case the edge sign is ambiguous in the TPS model at the protein level because some YHR131C phosphosites exhibit a significant increase in phosphorylation while others decrease. One YHR131C phosphosite is dependent on Rck2 during osmotic stress (Romanov et al., 2017), supporting our Rck2 –> YHR131C prediction. Similarly, we verified that 67 out of 91 (74%) predicted Cdc28 targets have at least one phosphosite with defective phosphorylation following Cdc28 inhibition (Holt et al., 2009; Kanshin et al., 2017) (Supplemental Results).

The high-quality TPS osmotic stress pathway demonstrates the algorithm is broadly useful beyond our own EGF stimulation study. It not only recovers many major elements of the classic HOG pathway representation but also prioritizes condition-specific kinase targets that are supported by independent perturbations.

## Discussion

The pathway structure illuminated by the phosphorylated proteins in our EGFR Flp-In cells differs considerably from the simple representations in pathway databases. Interpreting signaling data requires the reconstruction of models specific to the cells, stimuli, and environment being studied. TPS combines condition-specific information, time series phosphoproteomic data and the source of stimulation, with generic PPI networks and optional prior knowledge to produce *custom pathway representations*. The predicted EGFR signaling network highlights alternative connections to classic EGFR pathway kinases and extends the pathway with interactions that are supported by prior knowledge in other contexts or kinase inhibition. Combining different constraints on pathway structure from PPI network topology and temporal information is computationally challenging, and we identify predictions that can be obtained only through joint reasoning with all available data.

### Role of novel tested EGFR pathway edges

The EGFR pathway provides a rich background against which the TPS predictions can be validated, while concurrently letting us evaluate its ability to uncover new biology in an extensively studied and clinically relevant pathway. EGFR initiates a variety of signaling cascades that regulate phenotypes such as cell differentiation, migration, proliferation, and survival (Citri and Yarden, 2006). Dysregulation of EGFR activity by overexpression, mutation, or other mechanisms leads to cellular disease, including cancer (Huang et al., 2009; Pines et al., 2010).

All three experimentally-tested pathway edges naturally extend traditional EGFR pathway representations. These interactions involve kinases that play recognized roles in EGF response and introduce connections to target proteins that are controlled by the kinases in the EGFR context. We know *a priori* from the PPI network that the proteins physically interact in some manner. TPS illuminates the interactions’ relevance to EGF response and the details of the interactions, that is, the directions and signs.

ABL2, a well-characterized tyrosine kinase also known as ARG, is involved in actin remodeling, cell motility, and EGFR endocytosis (Colicelli, 2010). Though not detected in our mass spectrometry data, this protein is included in the TPS pathway due to its predicted role as a regulator of CRK, which is highly phosphorylated in response to EGF. ABL2 has been shown to phosphorylate CRK on Y221 *in vivo* (Wang et al., 1996). We confirm this site is sensitive to ABL2 inhibition, but it differs from the CRK site detected in our mass spectrometry data (Y136). KEGG’s ErbB signaling pathway depicts CRK as a direct regulator of ABL2, which contradicts our evidence that ABL2 directly phosphorylates CRK in the EGFR pathway. Both directions are potentially valid, but the current version of TPS does not model cycles, as discussed below.

The kinase MAPK1 (ERK2) is a central component of many stimulus responses and other biological processes whose dysfunction is linked with numerous diseases (Wortzel and Seger, 2011). ATP1A1 is the catalytic subunit of Na^+^,K^+^-ATPase, an ion pump that also plays a role in signal transduction when inhibited by ouabain (Reinhard et al., 2013), making it a candidate therapeutic target for diseases such as medulloblastoma (Wolle et al., 2014). MAPK1 phosphorylates ATP1A1 in response to insulin (Al-Khalili et al., 2004) and proinsulin-connecting peptide (C-peptide) (Zhong et al., 2004) stimulation, potentially on T81. However, both the TPS pathway model and our MAPK1 inhibition support an inhibitory effect on ATP1A1 phosphorylation. The difference could be attributed to the specific phosphorylation sites because MAPK1 was not linked to ATP1A1 tyrosine phosphorylation in other conditions (Al-Khalili et al., 2004). ATP1A1 Y55 (insignificant change) and Y260 (significant change) exhibit decreased phosphorylation in our mass spectrometry data. Our Western blot quantifies phosphorylation of Y10, which was detected jointly with Y260 phosphorylation in many PhosphoSitePlus Cell Signaling Technology curation sets (Hornbeck et al., 2015). Despite prior evidence of the MAPK1-ATP1A1 interaction, we cannot rule out indirect effects via other inhibited proteins (SCH772984 also impacts MAPK3) or an intermediate tyrosine phosphatase controlled by MAPK1, which could explain the significant change in ATP1A1 phosphorylation at 0 min (Figure 6D). Additional experiments are required to explore the mechanistic details of the EGF-induced relationships between these two proteins.

AKT1 is an important kinase in PI3 kinase signaling with roles in glucose transport, cell survival, cell growth, metabolism, and multiple diseases (Hers et al., 2011). Like ABL2, AKT1 phosphorylation was not detected in our mass spectrometry data, but it is correctly nominated as a regulator of other EGF-responsive proteins. AKT1 directly phosphorylates Zyxin on S142 (Chan et al., 2007), an activating interaction. Our Western blots support AKT1 inhibiting Zyxin phosphorylation based on an antibody specific to S142 and S143. The potential disagreement in edge sign is an intriguing topic for further study. It may suggest that although AKT1 directly phosphorylates Zyxin some contexts, their relationship in the EGF response context is indirect.

### Tradeoffs between ambiguity, expressiveness, and correctness

The modeling assumptions made when interpreting and translating biological data into logical constraints have complex effects on the degree of ambiguity, expressiveness, and accuracy of the resulting pathway summary. Even with temporal information, many pathway structures can explain the ordered signaling events. This motivates the reduction of ambiguity with hard logical constraints, where each constraint is fully trusted, instead of with probabilistic constraints (Hinton et al., 2006; Katoen et al., 2005), where a constraint can potentially be violated.

In the PPI network, we allow paths only through chains of experimentally detected PPI. In settings where the PPI network is less complete, we could include edges among highly correlated phosphorylated proteins or predicted interactions based on protein sequence, protein structure, pathway connectivity, or literature mining (Lees et al., 2011; Mosca et al., 2013). The pre-processing step that filters the PPI network operates on a weighted network. These additional edges could be assigned lower weights so that PCSF includes them in the TPS input network only if they are critical for connecting significantly phosphorylated proteins. This would reduce the impact of missing interactions on TPS pathways at the cost of potentially increasing ambiguity because there would be more possible paths through which signal can flow.

Likewise, we observe that some proteins, such as RAS and RAF family members, are not included in the TPS pathway because our mass spectrometry data do not detect their phosphorylation. To increase robustness to potential false negatives in the mass spectrometry, the input PPI network could be modified to include edges from relevant reference pathways with high weights (similar to (Patil et al., 2013)) so that PCSF prefers to include these interactions instead of other high-confidence connections in the PPI network. The weight of these prior knowledge edges would control the tradeoff between condition-specific *de novo* pathway discovery and conformance with prior knowledge.

Unlike single-cell mass cytometry data, where the peak activity times of a small number of phosphoproteins can be resolved precisely (Krishnaswamy et al., 2014), phosphorylation timing in cell population-level mass spectrometry data is inherently ambiguous. Therefore, instead of rigidly determining a protein’s time of activity by selecting the time point at which the greatest phosphorylation change is observed, TPS takes a more general approach. It allows a protein to be activated or inhibited whenever the phosphorylation significantly differs from the level before stimulation or at the immediately preceding time point as long as it is the *first time* at which that phosphorylation level has been observed. We focus on the initial pulse of signaling activity following stimulation, sampling more early time points in our EGF response study because we are more confident that these changes in phosphorylation intensity are due to PTMs instead of changes in protein abundance. Feedback loops cannot be detected when learning a single activation or inhibition time per peptide, a modeling decision we made in our three case studies. However, the TPS framework makes it possible to allow multiple activity changes per peptide in future applications. Statistical tests of the temporal phosphorylation profiles could determine the number of significant activity changes for each peptide. Then, TPS could search for pathway structures with feedback loops that explain the multiple activation or inhibition events per peptide. We describe a constraint-based solution that extends TPS to infer feedback in networks in Supplemental Experimental Procedures.

Lastly, we recognize that different phosphopeptides on the same protein can have different phosphorylation changes over time, and we allow each peptide to have its own activation times instead of forcing a single time per protein. This decision can lead to ambiguous edge direction predictions at the protein-level even when the directions are consistent at the peptide level. For example, DOCK1 interacts only with BCAR1 (Figure 5B), yet the direction and sign of the interaction are ambiguous. The uncertainty arises because BCAR1 is phosphorylated on both Y249 and Y387. TPS correctly concludes that the sign cannot be determined because one site could activate DOCK1 and then feed back and affect the other BCAR1 site.

### Contrasting TPS with related computational approaches

TPS provides a new way to integrate information from PPI networks, time series phosphoproteomic data, and prior knowledge by introducing a powerful constraint-based approach to build on concepts previously explored by related algorithms. Approaches for building networks from gene expression data alone (reviewed in (De Smet and Marchal, 2010)) can be applied to phosphoproteomic data as well. Extensions of these methods for temporal data introduce time lags and search for dependencies between genes’ expression levels over time (Zoppoli et al., 2010). Methods based on Granger causality (Masnadi-Shirazi et al., 2014) identify proteins whose phosphorylation predicts behavior at later time points and provide one type of causal model. However, as we showed in our comparison with the dynamic Bayesian network (Hill et al., 2012) and FunChisq (Zhang and Song, 2013), all methods that rely on the phosphorylation data alone (Henriques et al., 2017) will miss critical signaling pathway interactions because not all pathway members have observed phosphorylation changes.

Algorithms based on gene and protein perturbations provide an alternative approach toward causal models. Transcriptional regulatory networks have been inferred from expression changes induced by gene knockouts and knockdowns (Anchang et al., 2009; Markowetz et al., 2007; Wang et al., 2014; Yeang et al., 2004). Likewise, signaling networks have been reconstructed by stimulating a pathway and perturbing signaling nodes with kinase inhibitors or RNA interference. Protein activities are observed with antibody-based assays, and pathways are recovered *de novo* (Ciaccio et al., 2015; Fröhlich et al., 2009; Kiani and Kaderali, 2014; Molinelli et al., 2013) or by adapting prior pathway knowledge (Morris et al., 2011). The PHONEMeS method is unique for its ability to handle large-scale phosphoproteomic perturbation data (Terfve et al., 2015).

The HPN-DREAM network inference challenge (Hill et al., 2016) spawned several new approaches for analyzing time series phosphoproteomic data in multiple biological contexts. Participants predicted signaling pathways from *in silico* time series data and temporal reverse phase protein array data for approximately 45 phosphoproteins in four breast cancer cell lines under various stimuli and inhibitor treatments. In contrast, TPS focuses on reconstructing signed, directed signaling networks from large-scale phosphoproteomic data. The TPS networks rely on physical protein-protein interactions and include proteins that are not observed in the mass spectrometry. PropheticGranger (Carlin, 2014), the top performer in the HPN-DREAM experimental task, demonstrated the importance of prior knowledge in network inference and modified the standard Granger causality approach to assess dependencies between observed proteins. Meanwhile, TPS uses time series information to globally reason about temporally consistent network models, ensuring that all paths in a network agree with time series data and considering the temporal activities of nodes that are not direct neighbors in a path. FunChisq (Zhang and Song, 2013), the top performer in the HPN-DREAM *in silico* task, did not perform well on our EGF response data (Table S12).

In our EGFR study, the TPS PPI subnetwork input is provided by PCSF, but other network algorithms can also connect phosphorylated proteins using PPI. A related algorithm interpolates between globally optimal (Steiner tree) and locally optimal (shortest path) connections to different proteins (Yosef et al., 2009), and this method has been applied to link functional signaling proteins derived from phosphoproteomics data (Rudolph et al., 2016). Many other approaches connect source and target proteins in a PPI network to identify pathways. ResponseNet (Yeger-Lotem et al., 2009) does so with a maximum flow formulation; SHORTEST (Silverbush and Sharan, 2014) and PathLinker (Ritz et al., 2016) use shortest paths; Maximum Edge Orientation (MEO) (Gitter et al., 2011) orients the undirected edges to produce short, directed paths. Integer programs can express complex optimization preferences with multi-stage objective functions when predicting source-target connections (Chasman et al., 2014; MacGilvray et al., 2017). The predicted networks from any of these methods can be used as input for temporal analysis with TPS.

Among methods that integrate dynamic data and PPI networks, TPS is unique in its ability to assess and summarize all possible pathway structures that are consistent with the input network and the temporal constraints. TPS also considers all possible temporal activations for each peptide instead of mapping proteins to temporal bins in advance like TimeXNet (Patil and Nakai, 2014; Patil et al., 2013). Similarly, Budak et al. use time point-specific PCSF networks to map proteins to times (Budak et al., 2015), and TimePath assigns genes to transcriptional phases based on gene expression timing (Jain et al., 2016), and Khodaverdian et al. explore theoretical properties of temporal Steiner trees (Khodaverdian et al., 2016). The Signaling and Dynamic Regulatory Events Miner (SDREM) models temporal gene expression to infer the timing of transcription factor activity, but the pathway discovery phase does not use any temporal information (Gitter and Bar-Joseph, 2013; Gitter et al., 2013). Vinayagam et al. used temporal phosphorylation to evaluate their predicted PPI directions but did not consider dynamics when making the predictions (Vinayagam et al., 2011). Time series data and interaction networks have also been combined for inferring protein complex dynamics (Park and Bader, 2012), pathway enrichment (Jo et al., 2016), and related problems reviewed in Przytycka et al. (Przytycka et al., 2010).

The key difference between our work and other declarative computational approaches is that TPS operates on networks that are several orders of magnitude larger and summarizes very large solution spaces defined by sparser and less precise experimental data. Model checking and symbolic reasoning have been used to verify properties of manually constructed biological models (Fisher and Piterman, 2014), complete partially specified pathways using perturbation data (Köksal et al., 2013), and synthesize gene regulatory networks directly from data (Dunn et al., 2014; Moignard et al., 2015) (reviewed in (Fisher et al., 2014)). In addition, other types of declarative approaches, such as integer programming (Budak et al., 2015; Chasman et al., 2014; Jain et al., 2016; Ourfali et al., 2007; Sharan and Karp, 2013; Silverbush and Sharan, 2014) and answer set programming (Guziolowski et al., 2013), have been applied to biological pathway analysis. The TPS model summarization strategy, which makes it applicable to comprehensive signaling networks containing more than a hundred thousand edges and phosphosites, sets it apart from these related methods (Supplemental Results and Figure S15).

### Future directions in pathway synthesis

TPS offers a powerful framework for combining multiple types of declarative constraints to generate condition-specific signaling pathways. The constraint-based approach can be extended to include many additional types of data. New types of constraints could be derived from high-level properties that proteins, interactions, or pathways must satisfy. Future versions of TPS could incorporate perturbation data that links kinase inhibition or deletion to phosphorylation changes that are far downstream from the kinase. For instance, both temporal (Kanshin et al., 2015) and kinase perturbation (MacGilvray et al., 2017; Romanov et al., 2017) phosphoproteomic data are available for the yeast osmotic stress response. Modeling multiple related conditions (e.g., different ligand stimuli and inhibitor perturbations) could allow TPS to learn not only the signs of interactions but also the logic employed when multiple incoming signals influence a protein. Finally, TPS could accommodate user-defined assumptions or heuristics about pathway properties, such as restrictions on pathway length. Such complex constraints cannot be readily included in existing optimization-based approaches like dynamic Bayesian networks or TimeXNet.

As proteomic technologies continue to improve in terms of depth of coverage (Sharma et al., 2014; Zhou et al., 2013) and temporal resolution (Humphrey et al., 2015; Kanshin et al., 2015; Reddy et al., 2016), the need to systematically interpret these data will likewise grow. TPS enables reasoning with temporal phosphorylation changes and physical protein interactions to define what drives the vast protein modifications that are not represented by existing knowledge in pathway databases.

## Experimental Procedures

### Temporal Pathway Synthesizer algorithm overview

As illustrated in Figure 1, our algorithm receives three types of input: a time series mass spectrometry phosphoproteomic analysis of a stimulus response, an undirected graph obtained by filtering a large PPI network to identify interactions that are relevant to the differentially phosphorylated proteins, and optional prior knowledge about interaction directions (for example, kinase-substrate relationships).

The undirected input graph is obtained through a static analysis in which the significantly changing proteins are overlaid on a network of physical protein interactions. A network algorithm recovers connections among the affected proteins, simultaneously removing interactions that do not form critical connections between these proteins and nominating hidden proteins that do, even if they are not themselves phosphorylated. The specific criteria used to select proteins and interactions vary based on the network algorithm. Here we use PCSF (Tuncbag et al., 2013), but we have also successfully applied ResponseNet (Yeger-Lotem et al., 2009), MEO (Gitter et al., 2011), and TimeXNet (Patil and Nakai, 2014; Patil et al., 2013) for this step.

Our method combines the input data to recover pathways embedded in the network that agree with the temporal data. TPS transforms the input into logical constraints that determine which pathway models can explain the observed phosphoproteomic data. Topological constraints stem from the filtered PPI network and require that phosphorylated proteins are connected to the source of stimulation, such as EGF, by a cascade of signaling events. These signaling events propagate along the edges of the filtered PPI network. Temporal constraints ensure that the order of the signaling events is consistent with the timing of the phosphorylation changes. If protein B is downstream of protein A on the pathway, B cannot be activated or inhibited before A. Lastly, prior knowledge constraints guarantee that if the direction or sign of an interaction is known in advance, the pathway may not contain the edge with the opposite direction or sign. Typically, many possible pathways meet all constraints, so TPS summarizes the entire collection of valid pathways and identifies interactions that are used with the same direction or sign across all models. A symbolic solver reasons with these logical constraints and produces the pathway summary without explicitly enumerating all possible pathway models.

To illustrate this process, consider a hypothetical signaling pathway that contains a receptor node A and six other downstream proteins that respond when A is stimulated (Figure 7A). We cannot directly observe the pathway structure but seek to infer it from the types of data shown in Figure 7B - 7D. The first input is time series mass spectrometry data measuring the response to stimulating the receptor (node A), which detects phosphorylation activity for six proteins. Node B is absent from the phosphorylation data because it is ubiquitinated, not phosphorylated, by A. The second input is an undirected graph, which reveals high-confidence protein-protein interactions. These are detected independently of the stimulation condition but filtered based on their presumed relevance to the responding proteins with an algorithm such as PCSF. By combining phosphorylation data with the PPI subnetwork, this topology can recover "hidden" components of the pathway that are not phosphorylated (node B). Finally, our method accepts prior knowledge of directed kinase-substrate or phosphatase-substrate interactions, such as the edge C->D. Each of these inputs can be used individually to restrict the space of plausible pathway models. However, reasoning about them jointly produces a greater number of unambiguous predictions than considering each resource separately.

**Figure 7.**
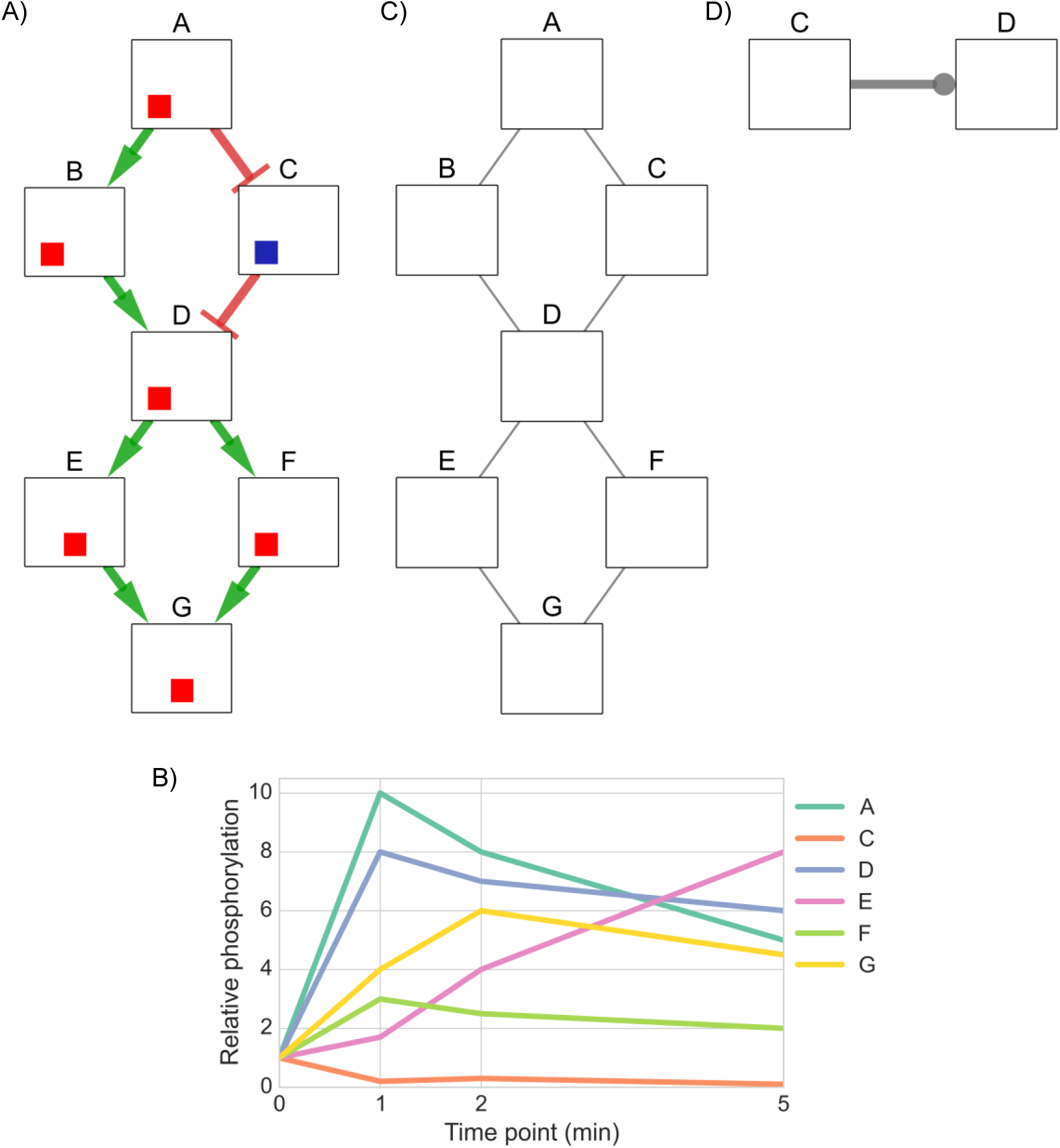
An artificial example illustrating the inputs to TPS. A) The signaling pathway that responds to stimulation of node A. The colored boxes on each node show the time at which the protein is activated or inhibited and begins influencing its downstream neighbor, with the leftmost position indicating the earlier time point. Red boxes are increases in activity, blue boxes are decreases, and white boxes are inactive time points, as explained in Figure 4B. The left position indicates the activity at 0 to 1 min, the center position at 1 to 2 min, and the right position at 2 to 5 min. B) The first input is time series phosphorylation data of the response to stimulating node A. C) The second input is an undirected graph of high-confidence interactions that can recover hidden components that do not appear in the temporal data, such as node B. D) The last input is prior knowledge of the pathway or the protein-protein interactions, expressed as (unsigned) directed edges. We represent unsigned edges with a circular arrowhead. Here, we have one such interaction, which is from C to D.

TPS exhaustively explores all signed, directed tree-structured pathway models, which are obtained by assigning signs and directions to edges of the undirected graph while restricting this space of networks through declarative constraints. These constraints are derived from the input. We next describe the constraints and how they restrict the space of models.

To formulate temporal constraints, we transform the time series data into a set of discrete signaling events (activation or inhibition) for each node, taking an event-based view of the signaling process (Table 1). We determine time points for each node that correspond to statistically significant phosphorylation changes. These discrete events are then used to rule out network models that contain signed, directed paths that violate the temporal ordering of these events no matter which event is chosen for each node. For example, there can be no edge from E to D in any model because D is activated strictly earlier than E regardless of whether E is activated at 1-2 min or 2-5 min. Because the time series data measures the response to a specific stimulus, we also devise topological constraints that ensure all signaling activity originates from this source. In our example, this asserts that all edges in a solution network must be on a directed path that starts at node A. Finally, our third input, the set of directed interactions, requires that no model violates this prior knowledge by including an edge from D to C.

**Table 1.** Plausible signaling events inferred for each node through a statistical analysis of the time series phosphorylation data. Although B is ubiquitinated in the 0-1 min interval, this is not observed in the phosphoproteomic input data.

| Node | Plausible temporal signaling events |
| --- | --- |
| A | Activated 0-1 min |
| B | Activated or inhibited at any time |
| C | Inhibited 0-1 min or 2-5 min |
| D | Activated 0-1 min |
| E | Activated 1-2 min or 2-5 min |
| F | Activated 0-1 min |
| G | Activated 0-1 min or 1-2 min |

We show in Figure 8 the pathway models that can be learned using each type of constraint alone and by asserting them jointly. When we enforce only temporal constraints, which corresponds to reasoning locally with phosphorylation data for pairs of nodes to see if one signaling event strictly precedes another, we obtain a single precise (signed and directed) prediction from D to E (Figure 8A). The topological constraints by themselves are sufficient to orient edges from the source A and from node D because D forms a bottleneck (Figure 8B). The prior knowledge constrains the direction of the edge from C to D, but its sign remains unknown (Figure 8C). Jointly enforcing all of these constraints has a nontrivial impact on the solution space (Figure 8D). For instance, we can infer that F must activate G. If the edge direction were reversed, F would be downstream of E, but the data show that activation of F precedes activation of E. The final model that includes all available data closely resembles the true pathway structure (Figure 7A). The edges incident to node B are ambiguous, and the interaction between E and G cannot be uniquely oriented, but all other interactions are recovered.

**Figure 8.**
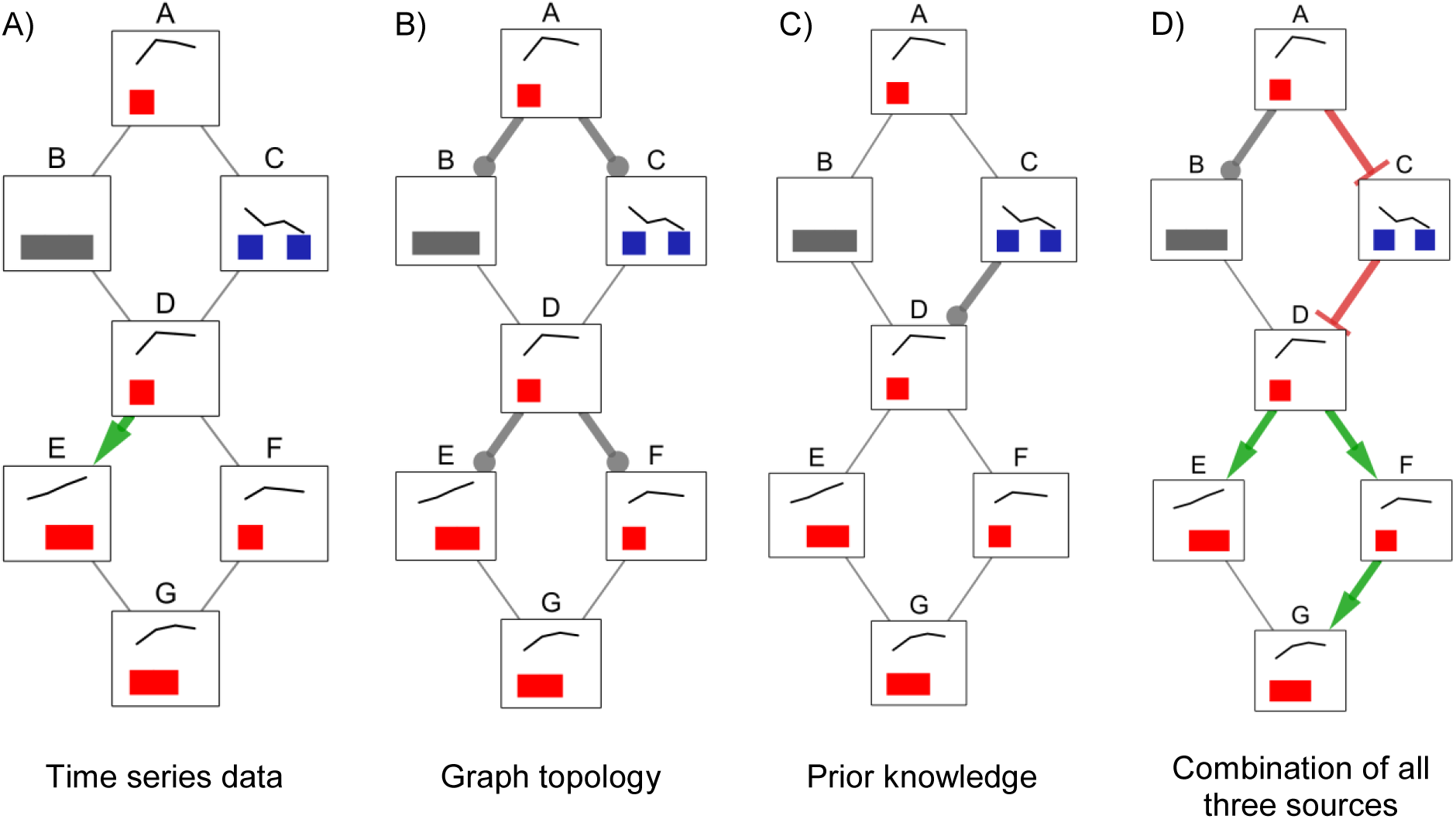
Summary graphs obtained by aggregating (via graph union) all possible signed, directed tree models for different constraints obtained from: A) time series data, B) graph topology, C) prior knowledge (in this example, kinase-substrate interaction directions), and D) all three types of input at the same time. If an edge has a unique sign and direction in a summary graph (colored green and red for activations and inhibitions, respectively), this means there are no valid models that assign a different orientation or sign to that edge. Edges that can have any combination of sign and direction in different models are gray without an arrowhead.

The summary for the combination of all constraints produces precise predictions that cannot be obtained by intersecting the summaries for the individual types of constraints. For instance, TPS infers that the relationship between F and G must be an activation from F to G because the sole way G can reach F in a tree rooted at A is through E, but F's activation precedes E's. This inference cannot be made by combining the models in panels A, B, and C. The simple example also highlights the differences in how the TPS constraint-based approach improves upon related methods based on correlation or the time point of maximum phosphorylation change (Supplemental Results and Figure S16).

### TPS pathway synthesis

TPS takes the undirected network of interactions produced by the PCSF algorithm and transforms it into a collection of signed, directed graphs that provide an explanation of dynamic signaling events.

#### Discretization of time series data

To find pathway models that agree with the phosphorylation dynamics, TPS first performs a discretization step that determines time intervals in which each protein may be differentially phosphorylated. The discrete set of activation and inhibition state changes is then used to rule out networks that violate the observed temporal behavior.

The transformation consists of finding time points for each profile where phosphorylation significantly differs from either the baseline (pre-stimulation) or the previous time point. In the baseline comparison, this time point is accepted only if it is not preceded by an earlier, larger change with respect to the baseline. If there is a hypothetical phosphorylation level at which the protein is activated and acts upon its downstream targets, a signaling event occurs only at the first time this threshold value is reached. This criterion does not apply when comparing to the phosphorylation level at the previous time point. In our EGF study, we use Tukey’s HSD test to find significant differential phosphorylation. If comparing a time point to the baseline or the previous measurement produces a p-value below a user-defined threshold, the time point is marked as a possible activation or inhibition event depending on whether the phosphorylation level increased or decreased relative to the earlier time point to which it was compared.

As an example, when we consider the profile for node E in Figure 7B, we find that both two and five minutes are time points where phosphorylation increases significantly relative to the previous time point (Table 1). As a result, both time points mark possible activation intervals. Even though the last measurement for node G significantly differs from the baseline, it does not constitute a possible activation because it is preceded by a larger value at 2 minutes. The hidden nodes for which there is no phosphorylation data (e.g., node B) are temporally unconstrained. They permit both activation and inhibition as possible state changes at all time intervals.

#### Modeling assumptions

Characteristics of the time series data directly influence our modeling assumptions. We assume at most one signaling event happens for every node across time points. Our logical solver can explore all possible activation and inhibition events for every node, but our experience shows that the data are too ambiguous to extend our interpretation beyond one event per node when modeling a single type of stimulation (such as EGF response). We also observe that, in the absence of perturbation experiments that test the pathway behavior under different initial conditions, it is impossible to distinguish between different Boolean logic functions governing the behavior of each node (AND/OR semantics) and whether a node exhibits activity in response to one or multiple predecessors. We therefore opt for signed, directed trees as our formalism for representing pathway models because they provide a sufficient basis for explaining the dynamic system behavior under these assumptions.

### Translating input into constraints

TPS transforms each input into a set of constraints that declaratively specify valid signed, directed tree models that agree with the data. These constraints are expressed as Boolean formulas with linear integer arithmetic, ranging over symbolic variables that represent choices on edge signs and orientations as well as how the temporal data are interpreted. The constraints can then be solved by a Satisfiability Modulo Theories (SMT) solver to find a network model that satisfies all constraints along with dynamic timing annotations for each interaction in the network.

Using constraints, we restrict the possible orientation and sign assignments to signed, directed tree networks rooted at the source node (e.g., EGF). Furthermore, constraints express how every tree model must agree with the time series data by establishing a correspondence between the order of nodes on tree paths and their temporal order of activity according to the time series data. Finally, we declaratively rule out models that contradict the prior knowledge of kinase-substrate interaction directions. In the Supplemental Experimental Procedures, we detail how such constraints are derived from the input. Together, these constraints typically define a very large space of candidate networks that agree with the data. TPS summarizes this space without explicitly enumerating all models.

### Pathway summaries

The space of all valid pathway models with timing annotations defined by the constraints we specified is typically very large, and enumerating all models is not computationally feasible. Given an undirected network G with V nodes and E edges, along with T time points, there are 5^E^ ways of assigning a sign and orientation to edges of G and (T*2 + 1)^V^ ways of assigning timing annotations to its nodes. Even for a network with 200 edges, the number of possible sign and orientation assignments is 6*10^139^. TPS can reason with even larger state spaces by producing summaries of all valid pathways instead of explicitly enumerating them.

We define a summary network as the graph union of all signed, directed tree networks that satisfy the stated constraints. Timing annotations are summarized by computing the set of possible annotations for each node over all solutions. Figure 8 shows an example of a pathway summary obtained by computing the union of all valid models in the solution space. In this union, we observe that some edges have a unique direction and sign combination, which signifies that this was the only observed signed, directed edge between two given edges across the solution space. However, this does not guarantee that the edge between the interacting proteins must be present in all valid pathway models. Meanwhile, when there are multiple direction and sign combinations between two nodes (e.g., between B and D), we know that multiple models have a different direction or sign assignment for the pair of nodes. The fourth summary graph indicates that at least two models contain an edge between B and D in opposite directions (Figure S17).

We compute the summary graph by performing a linear number of SMT solver queries in terms of the size of the input graph. Each query asks whether at least one signed, directed model contains a specific signed, directed edge. These individual queries are relatively computationally cheap in practice, and we can therefore have a view of the entire solution space without enumerating all models, which is typically intractable. The summary graph over-approximates the solution space. It is not possible to recover the exact set of valid models from the summary, but only a superset of the models (Supplemental Experimental Procedures and Figure S17). This tradeoff must be made in order to analyze such a large state space.

### Using solvers for synthesis

TPS uses the Z3 theorem prover (De Moura and Bjørner, 2008) via the ScalaZ3 interface (Köksal et al., 2011) to solve the constraints it generates. It additionally provides a custom solver implemented specifically for computing pathway summaries based on data-flow analysis. The custom solver and the symbolic solver produce identical pathway summaries. However, the custom solver is much more scalable because it is specifically designed to address our synthesis task, and can handle networks containing more than a hundred thousand edges and phosphosites (Supplemental Results and Figure S15).

### Cell culture, stimulation and generation of peptides

Flp-In 293 cells expressing EGFR were described previously (Gordus et al., 2009). These isogenic cells do not express EGFR heterodimerization receptor partners, and receptor quantities are uniform across cells (~100,000 EGFR/cell). Although their signaling response may differ from *in vivo* responses in human tissues, this system ensures the phosphorylation changes are EGFR-specific and reproducible across replicates. The cells were grown using standard cell culture procedures in Dulbecco's modified Eagle's medium supplemented with 10% (v/v) fetal bovine serum, 2 mM glutamine, 100 U/ml penicillin, 100 μg/ml streptomycin, and 150 μg/ml hygromycin B. For the activation of the EGF receptor, the cells were grown in plates to approximately 70% confluency, then washed once with phosphate-buffered saline (PBS) and incubated for 16 hours in serum-free medium. Subsequently cells were stimulated with 23.6 nM EGF (Peprotech) for 0, 2, 4, 8, 16, 32, 64, or 128 minutes. Untreated plates were used for the 0 min time point. After EGF stimulation, cells were lysed on ice with 3 ml of 8 M urea supplemented with 1 mM Na_3_VO_4_. A 10 μl aliquot was taken from each sample to perform the micro bicinchoninic acid protein concentration assay (Pierce) according to the manufacturer’s protocol. Cell lysates were reduced with 10 mM DTT for 1 hr at 56 °C, alkylated with 55 mM iodoacetamide for 45 min at room temperature, and diluted to 12 ml with 100 mM ammonium acetate; pH 8.9. 40 μg trypsin (Promega) was added to each sample (~200:1 substrate to trypsin ratio) and the lysates were digested overnight at room temperature. The whole cell digest solutions were acidified to pH 3 with acetic acid (HOAc) and loaded onto C18 Sep-Pak Plus C18 Cartridges (Waters). The peptides were desalted (10 mL 0.1% trifluoroacetic acid (TFA)) and eluted with 10 mL of a solution comprised of 40% acetonitrile (MeCN) with 0.1% TFA. Each sample was divided into ten aliquots and lyophilized overnight to dryness for storage at –80°C.

The peptides were then labeled using 8-plex iTRAQ reagents (Ab Sciex) according to the manufacturer’s instructions. 200 µg of lyophilized peptides were resuspended in 30 µl of dissolution buffer, and the corresponding iTRAQ reagent dissolved in 70 µl isopropanol was added. The mixtures were incubated at room temperature for 1 hr and concentrated to ~30 μl. Samples labeled with eight different isotopic iTRAQ reagents were combined and dried to completion. The sample was then rehydrated in 500 µl (0.1% HOAc) and desalted using a Sep-Pack Vac C18 column (Waters). The peptides were eluted with 80% acetonitrile, 0.1% HOAc. The eluate was evaporated to 100 µl in the SpeedVac and lyophilized.

### Phosphopeptide enrichment, mass spectrometry, and data analysis

Peptides containing phosphotyrosines were enriched using immunoprecipitation (IP). 12 µg of each of the antibodies P-Tyr-1000 (Cell Signaling Technologies), 4G10 (Millipore), and PT-66 (Sigma-Aldrich) were bound to 20 µl of packed protein G Plus agarose beads (Calbiochem) in IP buffer (100 mM Tris-HCl pH 7.4, 0.3% NP-40). The lyophilized peptides were rehydrated with IP buffer, and the pH of the solution was adjusted to 7.4 using 100 mM Tris-HCL ph 8.5. The peptide sample was added to the beads and incubated for 4 hours. The supernatant was then removed and saved for the next step. The beads were washed extensively using IP buffer, 100 mM Tris-HCl and H_2_O. The bound peptides were eluted using 50 µl 15% acetonitrile/0.1% TFA.

Serine and threonine (and remaining tyrosine) phosphorylated peptides were enriched from the IP supernatant using immobilized metal affinity chromatography. Briefly, protein concentration was adjusted to 1 mg/ml protein using wash buffer (80% MeCN/0.1%TFA). 100 µl of Ni-depleted Ni-NTA Superflow beads (Qiagen) were activated with 100 mM FeCl3. The supernatant was loaded onto the beads and incubated for 1hr. After washing the beads three times with wash buffer, the bound peptides were eluted twice with 1.4% ammonium hydroxide. The eluates were combined and evaporated in a SpeedVac to 5-10 µl. The sample was then reconstituted to a total volume of 20 µl in 20 mM ammonium formate pH 9.8.

All mass spectrometry experiments were performed on a Thermo Fisher Velos Orbitrap mass spectrometer equipped with a nanospray ionsource coupled to a nanoACQUITY Ultra Performance LC system (Waters) equipped with two binary pumps. Samples were separated using either 1D (IP eluate) or 2D (IMAC eluate) chromatography. For the 1D separation, the sample was loaded onto a 5 cm self-packed (Reliasil, 5 µm C18, Orochem) pre-column (inner diameter 150 μm) connected to a 20-cm self-packed (ReproSil, 3 µm C18, Dr. Maisch) analytical capillary column (inner diameter 50 μm) with an integrated electrospray tip (∼1 μm orifice). Peptides were separated using a 115-minute gradient with solvents A (H_2_O/formic acid (FA), 99.9:1 (v/v)) and B (MeCN/FA, 99.9:1 (v/v)) as follows: 1 min at 2% B, 84 min from 98 to 40% B, 5 min at 40% B, 20 min at 20% B, and 14 min at 2% B. For the 2D reverse phase chromatography, the sample was first loaded onto a 5 cm self-packed Xbridge column (Waters, inner diameter 150 μm) and eluted with a 7-step gradient of 1, 3, 6, 9, 13, 25, and 44% B with solvents A (H_2_O/20 mM ammonium formate pH 9.8) and B (MeCN/20 mM ammonium formate, pH 9.8). The eluted sample was directly loaded onto a 5 cm self-packed precolumn (Reliasil, 5 µm C18, Orochem), which was connected to a 20 cm self-packed analytical column (ReproSil, 3 µm C18, Dr. Maisch). The peptides were eluted with the same gradient as described above in the second dimension.

Eluted peptides were directly analyzed using a Velos-Orbitrap mass spectrometer operated in data-dependent acquisition (DDA) mode to automatically switch between MS and MS/MS acquisitions. The Top 10 method was used, in which full-scan MS (from m/z 350–2000) was acquired in the Orbitrap analyzer at 120,000 resolution, followed by high-energy, collision-induced dissociation (HCD) MS/MS analysis (from m/z 100–1700) of the top 10 most intense precursor ions with a charge state >2. The HCD MS/MS scans were acquired using the Orbitrap analyzer at 15,000 resolution at a normalized collision energy of 45%, with the ion selection threshold set to 10,000 counts. Precursor ion isolation width of 2 m/z was used for the MS/MS scans, and the maximum allowed ion accumulation times were set to 500 ms for full (MS) scans and 250 ms for HCD (MS/MS). The standard mass spectrometer tune method settings were as follows: Spray Voltage, 2.2 kV; no sheath and auxiliary gas flow; heated capillary temperature, 325 C; Automatic Gain Control (AGC) enabled. All samples were analyzed by LC-MS/MS in biological triplicates.

MS/MS data files were searched against the human protein database using Comet (Eng et al., 2013). Variable (phosphorylation of serine, threonine, or tyrosine, 79.966331 Da, methionine oxidation, 15.9949 Da) and static (carbamidomethylation of cysteine, 57.02 Da and the iTRAQ modification of 304.205360 Da to peptide N-terminus and lysine side-chains) modifications were used for the search (Supplemental File 9). We applied a 1% false discovery rate threshold at the peptide level because our analysis models individual phosphopeptides. Quantitation of the iTRAQ signals was performed using Libra (Deutsch et al., 2010) (see Supplemental Experimental Procedures for pre-processing and normalization).

Across the three biological replicates, we quantified 5442 unique peptides in at least one replicate and 1,068 peptides in all replicates. We focused on these 1,068 peptides for the computational modeling because the repeated observations indicate more reliable quantification and let us assess the significance of phosphorylation changes (Table S3). The Supplemental Experimental Procedures describe the Tukey's Honest Significant Difference statistical testing (Yandell, 1997) and temporal phosphorylation analysis. Table S14 demonstrates robustness to the p-value threshold.

### Quantitative Western blotting

The following kinase inhibitors were used at the following concentrations: 25 nM Dasatinib (#S1021), 400 nM SCH772984 (#S7101), and 800 nM MK-2206 (#S1078, all Selleckchem). The Flp-In 293 EGFR cells were serum-starved for 16 hours in growth medium without FBS. Then, if indicated, the kinase inhibitors were added and incubated with the cells for 60 min. Only DMSO was added to control cells. After stimulation with 23.6 nM EGF for the indicated times, the cells were lysed using RIPA buffer (25mM Tris-HCl pH 7.6, 150mM NaCl, 1% NP-40, 1% sodium deoxycholate, 0.1% SDS) with 1 mM sodium orthovanadate. Cells without EGF stimulation were used as controls (0 min time point). After incubation on ice for 15 minutes, the lysates were centrifuged at 21000 g for 10 min, and the protein concentration of the supernatants was determined. Protein amounts were adjusted to 30–50 ug protein/well, and protein phosphorylation was assessed by SDS separation and Western blotting. The membranes were probed with the following antibodies at a dilution of 1:1000: pY221-CRK (#3491, Crk-II isoform), pY10-ATP1A1 (#3060), and pS142/143-Zyxin (#8467, all Cell Signaling Technologies). β-actin (#3700) was used to normalize loading across the gel. Fluorescently labeled secondary antibodies were added according to the manufacturer’s instructions at 1:5000 (Goat anti rabbit IRDye 680 and Goat anti mouse IRDye 800, Li-COR Biosciences). Blots were imaged using an Odyssey Infrared Imaging System (Li-COR Biosciences). Quantification of the phosphorylated proteins was performed with the Odyssey analysis software.

### Prize-collecting Steiner forest

We use the prize-collecting Steiner forest implementation from Omics Integrator (Tuncbag et al., 2016) to recover the most relevant PPIs connecting the phosphorylated proteins. PCSF recovers the sparse subnetwork *F* = (*V*_*F*_,*E*_*F*_) from a dense PPI network that links proteins of interest by solving

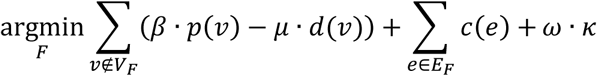

where *p* is a positive score (prize) that reflects the relevance of a vertex (protein) *v*, *d* is the degree (number of neighbors) of *v*, *c* is a positive cost for including an edge (interaction) *e* in the subnetwork, and κ is the number of disconnected trees in the subnetwork. Parameters β, μ, and ω control the size and structure of the solution subnetwork. An advantage of PCSF is that it nominates pathway members that are not detected by the mass spectrometry but form critical pathway connections to phosphorylated proteins, like ABL2 and AKT1 in our EGF response study (Figure 5). The Supplemental Experimental Procedures describe how we set these parameters, ran PCSF multiple times to identify parallel connections between proteins, generated prizes from the phosphoproteomic data, and created a weighted interaction network from iRefIndex (Razick et al., 2008) and PhosphoSitePlus (Hornbeck et al., 2015) PPI. Omics Integrator solves the PCSF optimization problem with the belief propagation-based msgsteiner algorithm (Bailly-Bechet et al., 2011).

### Data and software availability

The raw mass spectrometry proteomics data have been deposited to the ProteomeXchange Consortium via the PRIDE (Vizcaíno et al., 2016) partner repository with the dataset identifier PXD006697, and the processed data are available in Tables S1 – S3. The latest version of TPS is available at https://github.com/koksal/tps as MIT-licensed open source software. TPS version 2.1 and instructions for running the software are included in Supplemental File 10. Our prototype visualization tool for TPS output is available at https://github.com/koksal/tpv as MIT-licensed open source software.

## Author contributions

Conceptualization, A.S.K., S.S., R.B., A.W.-Y., E.F., J.F., and A.G.; Methodology, A.S.K., K.B., R.B., A.W.-Y., E.F., J.F., and A.G.; Software, A.S.K. and A.G.; Validation, K.B. and N.D.C.; Formal Analysis, A.S.K., D.R.C., and A.G.; Investigation, A.S.K., K.B., D.R.C., A.M., N.D.C., M.E.M., and A.G.; Data Curation, A.S.K., K.B., D.R.C., A.W.-Y., and A.G.; Writing, A.S.K., K.B., M.E.M., R.B., A.W.-Y., E.F., J.F., and A.G.; Visualization, A.S.K., E.F., and A.G.; Supervision, R.B., A.W.-Y., E.F., J.F., and A.G.

## Acknowledgements

We thank Nir Piterman for algorithmic discussions, Anthony Soltis and Jennifer Wilson for assistance preparing the human protein-protein interaction network, Alex Hu for phosphoproteomic data processing, Evgeny Kanshin for assistance with the yeast phosphoproteomic data, Audrey Gasch for yeast osmotic stress response discussions, and Sandra Kaplan for copy editing. Microsoft Research supported A.S.K. and A.G. during the initial stages of this work. A.G. acknowledges support from NSF grant DBI-1553206. D.R.C. acknowledges support from NIH grant U54-AI117924. M.E.M. acknowledges support from NIH training grant T32-HG002760. E.F. acknowledges support from NIH grant U54-NS091046. R.B. acknowledges support from NSF grants CCF-1139138, CCF-1337415, and ACI-1535191, a grant from the U.S. Department of Energy, Office of Science, Office of Basic Energy Sciences Energy Frontier Research Centers program under award number FOA-0000619, and grants from DARPA FA8750-14-C-0011 and DARPA FA8750-16-2-0032 as well as gifts from Google, Intel, Mozilla, Nokia, and Qualcomm.

## Supplemental File Index

- **Table S1.** Unprocessed mass spectrometry data. Three biological replicates with two technical replicates each. IP and IMAC are listed separately.
- **Table S2.** Normalized mass spectrometry data for each of the biological replicates.
- **Table S3.** Processed mass spectrometry data for peptides that appear in all biological replicates. Includes median intensity at each time point, log2 fold change, and statistical significance from Tukey’s HSD test.
- **Table S4.** All interactions in the TPS pathway for our EGF response data. Includes the directions and signs each interaction may take and the evaluation results with respect to EGFR reference pathways, kinase-substrate interactions, and NLP.
- **Table S12.** The overlap between the TPS, TimeXNet, FunChisq, and DBN pathway predictions and reference pathways.
- **Supplemental File 1.** Mass spectrometry data formatted as input for the PCSF and TPS algorithms.
- **Supplemental File 2.** Subnetwork output by PCSF run on our EGF response data as a Simple Interaction Format (SIF) file.
- **Supplemental File 3.** Cytoscape (Shannon et al., 2003) session file for visualizing the TPS pathway for our EGF response data. Created with Cytoscape version 3.2.0.
- **Supplemental File 4.** Web pages containing a detailed evaluation of TPS predictions with respect to reference EGFR pathways and kinase-substrate interactions.
- **Supplemental File 5.** Subnetwork output by PCSF run on the Olsen et al. EGF response data as a SIF file.
- **Supplemental File 6.** Cytoscape session file for visualizing the TPS pathway for the Olsen et al. EGF response data.
- **Supplemental File 7.** Subnetwork output by PCSF run on the yeast osmotic stress response data as a SIF file.
- **Supplemental File 8.** Cytoscape session file for visualizing the TPS pathway for the yeast osmotic stress response data.
- **Supplemental File 9.** Comet MS/MS search engine parameters file.
- **Supplemental File 10.** Version 2.1 of the TPS code, including instructions for running the software, example data, and scripts for linking PCSF and TPS. The most recent version of the code is available at https://github.com/koksal/tps.

